# Multiple time scales of adaptation allow cones to encode the inputs created by visual exploration of natural scenes

**DOI:** 10.1101/2021.02.13.431101

**Authors:** Juan M. Angueyra, Jacob Baudin, Gregory W. Schwartz, Fred Rieke

## Abstract

Primates explore their visual environment by making frequent *saccades*, discrete and ballistic eye movements that direct the fovea to specific regions of interest. Saccades produce large and rapid changes in input. The magnitude of these changes and the limited signaling range of visual neurons means that effective encoding requires rapid adaptation. Here, we explore how cone photoreceptors maintain sensitivity under these conditions. Adaptation makes cone responses to naturalistic stimuli highly nonlinear and dependent on stimulus history. Such responses cannot be explained by linear or linear-nonlinear models but are well explained by a biophysical model of phototransduction with fast and slow adaptational mechanisms. The resulting model can predict cone responses to a broad range of stimuli and enables the design of stimuli that elicit specific (e.g. linear) cone photocurrents. These advances will provide a foundation for investigating the contributions of cones and post-cone processing to visual function.

## INTRODUCTION

Everyday visual activities, like reading or identifying familiar faces in a crowd, rely on signaling in the fovea, a small and specialized region of the retina where cone photoreceptor density and perceptual spatial acuity are highest (reviewed by (Rodieck, 1973)). Most visual information is encoded during *fixations*, periods of time where gaze is relatively stationary on the visual scene. Visual cues detected in the periphery, where spatial acuity and cone density are lower, cause ballistic eye movements, or *saccades*, that direct the fovea to the region of interest. Humans typically make multiple saccades every second, and each saccade can span several degrees of visual angle (Harris, Hainline, Abramov, Lemerise, & Camenzuli, 1988). On the spatial scale of saccades, natural scenes can exhibit large differences in local intensity and local spatial contrast — i.e. the fluctuations in intensity about the mean in small image patches (Frazor & Geisler, 2006).

Several issues make reliably encoding the visual inputs encountered during eye movements challenging. First, given that the dynamic range of neural signals is small compared to the range of inputs encountered during different fixations, the visual system must adaptively adjust sensitivity to match the prevailing inputs. Such adaptation must occur locally in the retina, given the large differences in inputs in different regions of a scene. Second, given that fixations only last 200–600 ms, adaptational mechanisms must operate quickly so as to match neural sensitivity to the inputs encountered within a fixation rather than those encountered over previous fixations.

The need to adapt to an ever-changing environment is ubiquitous across sensory systems. For example, adaptation allows bacteria to follow molecular gradients across a > 10,000-fold range of concentrations (Bialek & Setayeshgar, 2005; Neumann et al., 2014), and the kinetics of adaptation govern the ability to follow these gradients (Block, Segall, & Berg, 1983). Similar challenges arise in the tracking of odor plumes in insects, where turbulent flow creates enormous variations in odorant concentrations (Carde & Willis, 2008), and in the auditory system, where behaviorally-relevant sounds span intensities that can differ by at least nine orders of magnitude (Viemeister & Bacon, 1988). In olfaction and audition, adaptation at the primary receptors (odorant receptor neurons and hair cells) is essential to maintain sensitivity (Kelliher, Ziesmann, Munger, Reed, & Zufall, 2003; Fettiplace & Ricci, 2003; Gorur-Shandilya, Demir, Long, Clark, & Emonet, 2017).

The primary visual receptors - rod and cone photoreceptors - also adapt strongly (reviewed by (Fain, 2001; Burns & Baylor, 2001)). Adaptation in photoreceptors affects both the gain and kinetics with which light inputs are converted to electrical signals. For typical daytime light levels, adaptation in the retinal output to changes in mean intensity is dominated by adaptation in the cones themselves (Dunn, Lankheet, & Rieke, 2007). We have a good understanding of how photoreceptor adaptation contributes to maintaining visual sensitivity to slowly changing inputs—e.g. the rising or setting sun— and of the mechanistic basis of photoreceptor adaptation, particularly in rods (reviewed by (Fain, 2001; Burns & Baylor, 2001)). We know much less about how photoreceptor adaptation contributes to reliable encoding of the large and rapid changes encountered as gaze shifts within a visual scene. Our focus here is on understanding the encoding of such naturalistic inputs by peripheral primate cones and using this understanding to construct models that allow us to predict and manipulate cone responses to a wide range of inputs. The ability to manipulate cone responses provides a needed tool to probe the causal role of cone signaling properties on responses in subsequent visual neurons and on behavior.

## RESULTS

The results are divided into four sections. First, we show that time-dependent adaptation strongly shapes the responses of peripheral primate cones to stimuli with large and rapid changes in intensity like those encountered during eye movements. Second, we characterize the kinetics of adaptation for a diverse set of stimuli. Third, we incorporate these measurements into a biophysical model able to account for cone responses across these stimuli. Fourth, we show two examples of how the model can be used to explore the role of cones in coding by downstream neurons.

### Primate cone responses to naturalistic stimuli are highly nonlinear

We start by describing responses to stimuli that approximate the intensity changes encountered by single cones during natural vision (Figure 1A). We ignored fixational eye movements (i.e. microsaccades, tremor and drift) and focused on saccades and fixations. We modeled the duration of fixations as an exponential distribution with a minimum interval between saccades (Harris et al., 1988) and a time constant that produced ∼3 saccades every second. The light intensity during each simulated fixation was determined by randomly sampling from an intensity distribution taken from natural images (van Hateren & Snippe, 2007). The intensity changed linearly from the value at one fixation to that at the next fixation over 15 ms (see Methods for details). The resulting stimuli capture the large and rapid changes in light intensity characteristic of the inputs that cones encounter during natural vision (Figure 1B, top).

**Figure 1.**
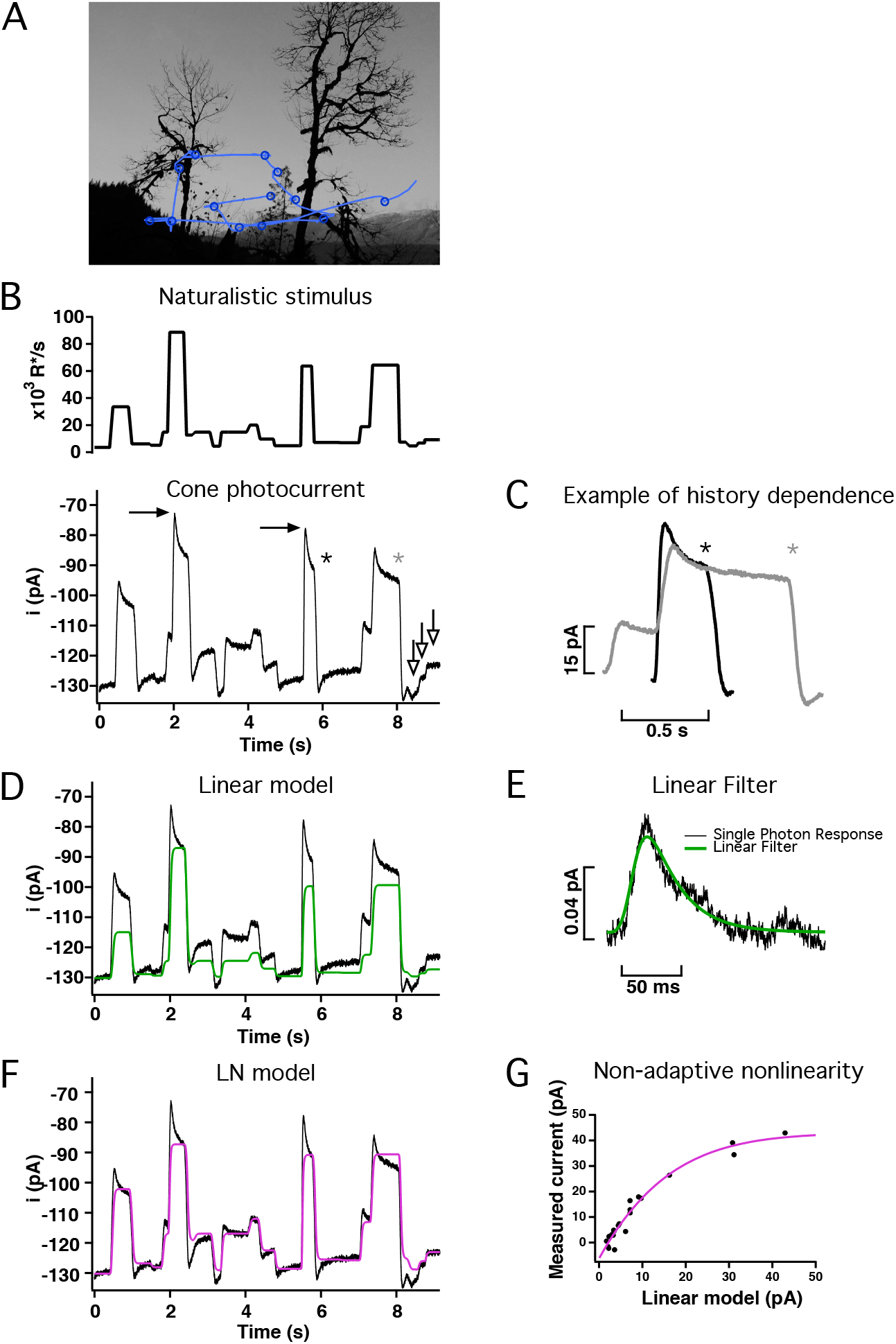
Responses of primate cones to naturalistic stimuli are not well captured by linear or linear-nonlinear (LN) models. **A**. Schematic of eye movements (blue lines) and fixations (blue circles) during free-viewing of a natural scene. **B**. (top) Stimulus emulating the large and frequent changes in mean light intensity experienced by a single cone during free-viewing. (bottom) Cone responses to this stimulus are highly nonlinear. For example, the difference between the responses marked by the filled black arrows is similar to the difference in responses marked by the open white arrows even though the corresponding stimulus intensities differ 10-fold. **C**. History dependence exemplified by two responses to the same light intensity but proceeded by different light intensities (asterisks in B). **D**. Linear model (green trace) scaled to match the final current at the highest light intensity. The model fails to accurately predict responses to most intensities and does not capture the response dynamics following a change in light intensity. **E**. Estimated single-photon response for cone in B. The fit to the response (see Methods) was used as a filter to construct a linear estimate of the response in D. **F**. A linear—nonlinear (or LN) model (magenta trace) captures the currents at the end of fixations but still fails to capture the dynamics of the response. **G**. The LN model was built using a non-adaptive nonlinearity constructed by fitting the relation between the measured currents (after baseline subtraction) at the end of fixations (y-axis) and the linear model (x-axis).

We delivered these naturalistic stimuli while recording current responses of voltage-clamped cones. These currents are dominated by phototransduction in the outer segment of the recorded cone and contain negligible contributions from electrically coupled cones or from horizontal-cell feedback (Dunn et al., 2007; Angueyra & Rieke, 2013). Cone current responses exhibited at least three signs of nonlinearity (Figure 1B, bottom). First, responses following increases or decreases in light intensity were not equal and opposite as would be expected for a linear system. In particular, increases in intensity elicited more pronounced current transients than decreases (e.g. the responses to the increase and the decrease in intensity between 5.5 and 6 s in Figure 1B). Second, gain was not constant. Instead, high intensities produced considerable response compression; for example, the difference in the responses produced by the two highest intensities in Figure 1B (filled arrows) were similar in magnitude to the difference in the responses produced by much smaller intensity changes when overall intensity was lower (open arrows). Third, the responses showed history dependence, such that responses to particular light intensities depended on the previous intensities. For example, steps to a common intensity from different starting intensities elicited responses that differed in both peak amplitude and kinetics (Figure 1B, asterisks and Figure 1C); such differences in kinetics would not be expected if responses were linear.

Not surprisingly, the nonlinear properties of the cone responses illustrated in Figure 1 could not be captured by linear models (Figure 1D and E) or by models that incorporate a non-adaptive (i.e. static/time-independent) nonlinearity (Figure 1F and G). Models for ganglion-cell responses often implicitly assume that early retinal processing, including the cone responses, are near-linear and that the dominant nonlinearities in the ganglion-cell responses originate in post-cone retinal circuits (this includes linear-nonlinear, stacked linear-nonlinear and generalized-linear models) (Chichilnisky, 2001; Pillow et al., 2008). Such models may benefit from incorporating time-dependent nonlinearities in the cones given the large impact of these nonlinearities on responses to the large and rapid changes encountered under natural conditions.

### Kinetics of adaptation

Time-dependent nonlinearities are pronounced in cones from many species (Soo, Detwiler, & Rieke, 2008; Korenbrot, 2012; Schnapf, Nunn, Meister, & Baylor, 1990; Schneeweis & Schnapf, 2000; Angueyra & Rieke, 2013; Cao, Luo, & Yau, 2014). Such nonlinearities are likely to be strongly engaged by naturalistic inputs, but their kinetics have not been well characterized for primate cones. The experiments described below characterize the time course of cone light adaptation using several types of stimuli. These results provide an important constraint for models of the cone response.

#### Primate cones exhibit fast and slow light adaptation

To determine the time course of adaptation, we probed how gain changes as a function of time following an abrupt increase or decrease in mean light level. We delivered brief flashes with variable delays relative to the onset and offset of a light step, and isolated the flash responses by subtracting the response to the step alone (Figure 2A-C). Flashes delivered prior to step onset or well after step offset elicited unadapted flash responses. A flash delivered near the end of the step elicited a completely light-adapted response. Flashes delivered at times near step onset or offset probed the transition between unadapted and adapted responses (Figure 2C).

**Figure 2.**
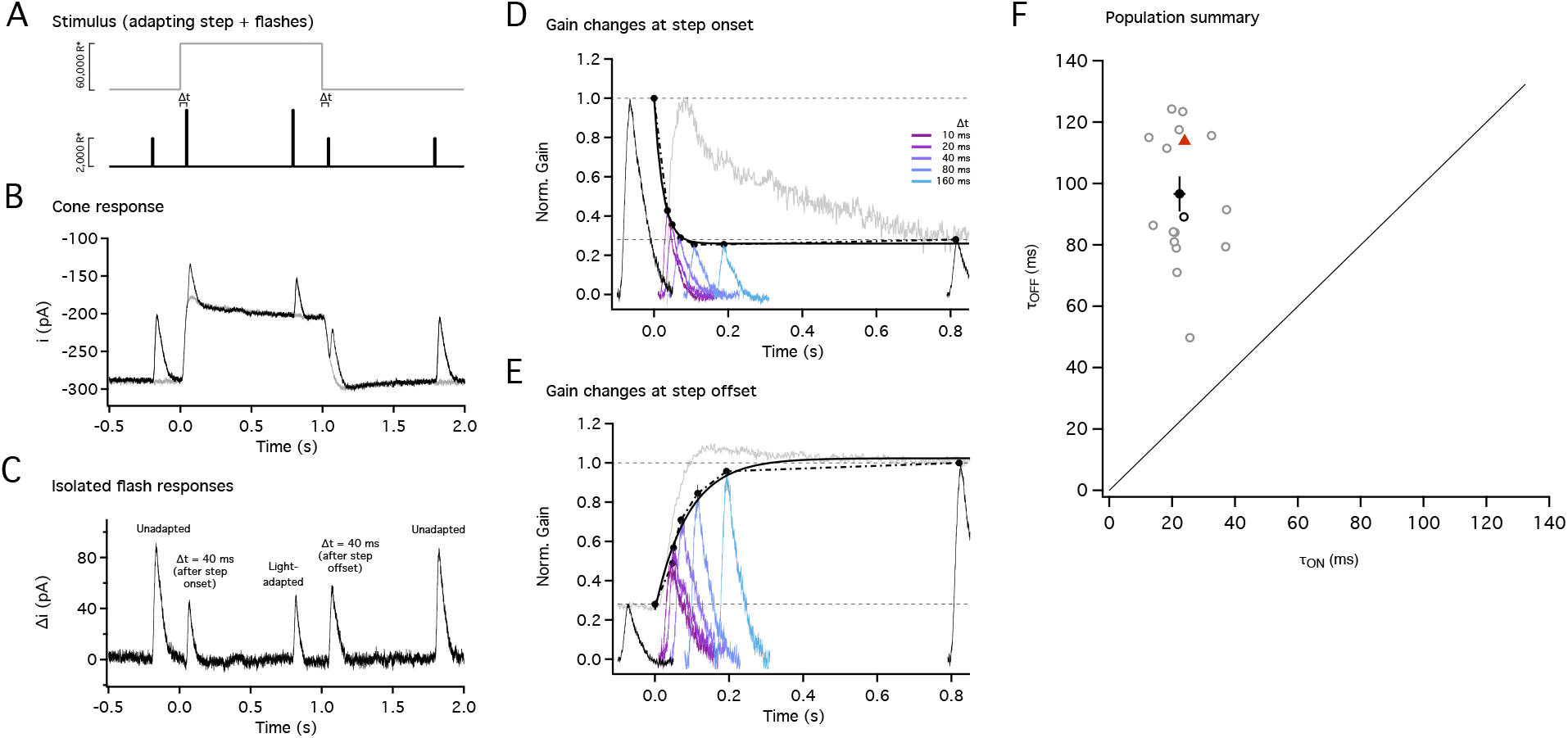
Gain changes during light-adaptation are fast and well-tuned to the duration of fixations. **A**. Stimulus used to probe the kinetics of gain changes during light adaptation. Five flashes (black trace) were superimposed on an adapting step (gray trace); the first, third and fifth flashes were fixed in time (black), while the second and fourth were delivered with variable delays (Δt) from step onset and offset. Flashes during the step were 2-fold brighter to partially counteract light adaptation. For this example trace, Δt = 40 ms. **B**. Average responses to the adapting step alone (gray trace) or in combination with the five flashes (black trace) for Δt = 40 ms. **C**. Flash responses isolated by subtracting the response to the step alone. The first and fifth flash produced unadapted responses, while the smaller and faster response to the third flash (near the end of the step) reflected adaptation. The flashes following step onset and offset elicited responses in transition between the two states. **D**. Gain changes rapidly at step onset. Gain measurements obtained by dividing the response by the flash strength and normalizing to the gain in darkness; black traces correspond to gain in darkness (leftmost trace) and to steady-state adapted gain (rightmost trace). Colored traces correspond to flashes with a variable delay (Δt) from the step onset. The speed of the gain changes was tracked by identifying the peaks and approximating their time course with an exponential function. The time constant of the best fit exponential was τ_On_ = 14 ms (black smooth line). **E**. Gain changes more slowly at step offset. Black traces correspond to steady-state adapted gain (leftmost trace) and gain in darkness (rightmost trace). Colored traces correspond to flashes with a variable delay from the step offset (same delays as in D). The time constant of the best fit exponential was τ_Off_ = 86 ms (black smooth line). For **D** and **E**, the response to the step without flashes has been displaced and rescaled to compare kinetics (gray traces). **F**. Collected time constants for gain changes at step onset and offset. Mean and SEM are shown as black circle and error bars, individual cells are shown as gray open circles (n = 15) and the example cell in (A-D) is shown as the black open circle. All cells lie above the unity line (black dashed line). The time constants for the biophysical model (see text and Figure 6) are shown by the red triangle.

The response gain was estimated by dividing the isolated flash responses by the flash strength. Changes in gain following both step onset and offset were largely complete within 200 ms — i.e. within the duration of a typical fixation between saccades; however, gain changes following step onset were faster than those following step offset (Figure 2D and E). Approximate time constants were extracted by fitting the gain changes with single exponential functions (black lines in Figure 2D and E). Across recorded cells (n = 15) and step intensities (1,500 - 100,000 R*/s), the extracted time constants of the gain changes were 3 to 4 times faster at step onset than at step offset (Figure 2F; mean ± SEM: τ_onset_ = 23 ± 2 ms; τ_offset_ = 94 ± 5 ms; p < 10^−8^ for τ_offset_ > τ_onset_). Adaptation following step onset sped with increasing light level, while that following step offset did not change significantly (Figure 2 - Figure Supplement 1).

The response to the step itself took ∼40 ms to reach peak and then decayed slowly to a maintained level (grey trace in Figure 2B). Most of the changes in flash-response gain occurred during the rising phase of the step response (grey trace in Figure 2D). A small *increase* in gain during the slow decay in the step response likely originated from the slow increase in circulating current (compare the amplitudes of the blue and purple flash responses to the response to the step itself in grey in Figure 2D). The current response to step offset exhibited two phases: an initial rapid recovery that overshot the baseline current, followed by a gradual return to baseline. Changes in gain persisted well beyond the rapid recovery phase and more closely followed the slow return to baseline (Figure 2B and E).

#### Kinetics of onset and offset of Weber adaptation

Adaptation in cones closely follows Weber’s law — i.e. across a broad range of light levels, gain is inversely related to mean light level (Burkhardt, 1994; Schneeweis & Schnapf, 2000; Dunn et al., 2007; Angueyra & Rieke, 2013). Weber’s law predicts that responses to stimuli with fixed contrast will be independent of mean light level. The experiments of Figure 2 suggest that adaptation occurs rapidly following a change in mean light level, and hence that Weber’s law should hold shortly after a change in light level. To test this prediction directly, we replaced the light flashes in Figure 2 with sinusoids of fixed contrast and explored steps from or to a common mean light level.

These experiments required long-lasting and stable recordings; hence we used perforated-patch recordings to avoid the washout of internal components that occurs during tight-seal whole-cell recordings; to avoid voltage-clamp errors associated with the higher access resistance in perforated-patch recordings, we measured photovoltages rather than photocurrents.

The onset of Weber adaptation was probed by recording responses to steps from low to high mean light levels with sinusoidal stimuli superimposed. Figure 3A shows an experiment in which we stepped from a single low light level to two different high light levels. We isolated responses to the sinusoidal stimuli by subtracting responses to the steps delivered alone (Figure 3B). If the cone response followed the stimulus veridically, responses to the sinusoidal stimuli should differ almost 2-fold at the two mean light levels. Instead, sinusoidal responses at the two different light steps were similar even shortly after the light step, indicating that contrast invariance was achieved quickly (Figure 3C and D). Indeed, responses exhibited contrast-invariance by the time at which the response to the light step reached its peak and well before the voltage sagged to reach its final steady-steady level (Figure 3C).

**Figure 3.**
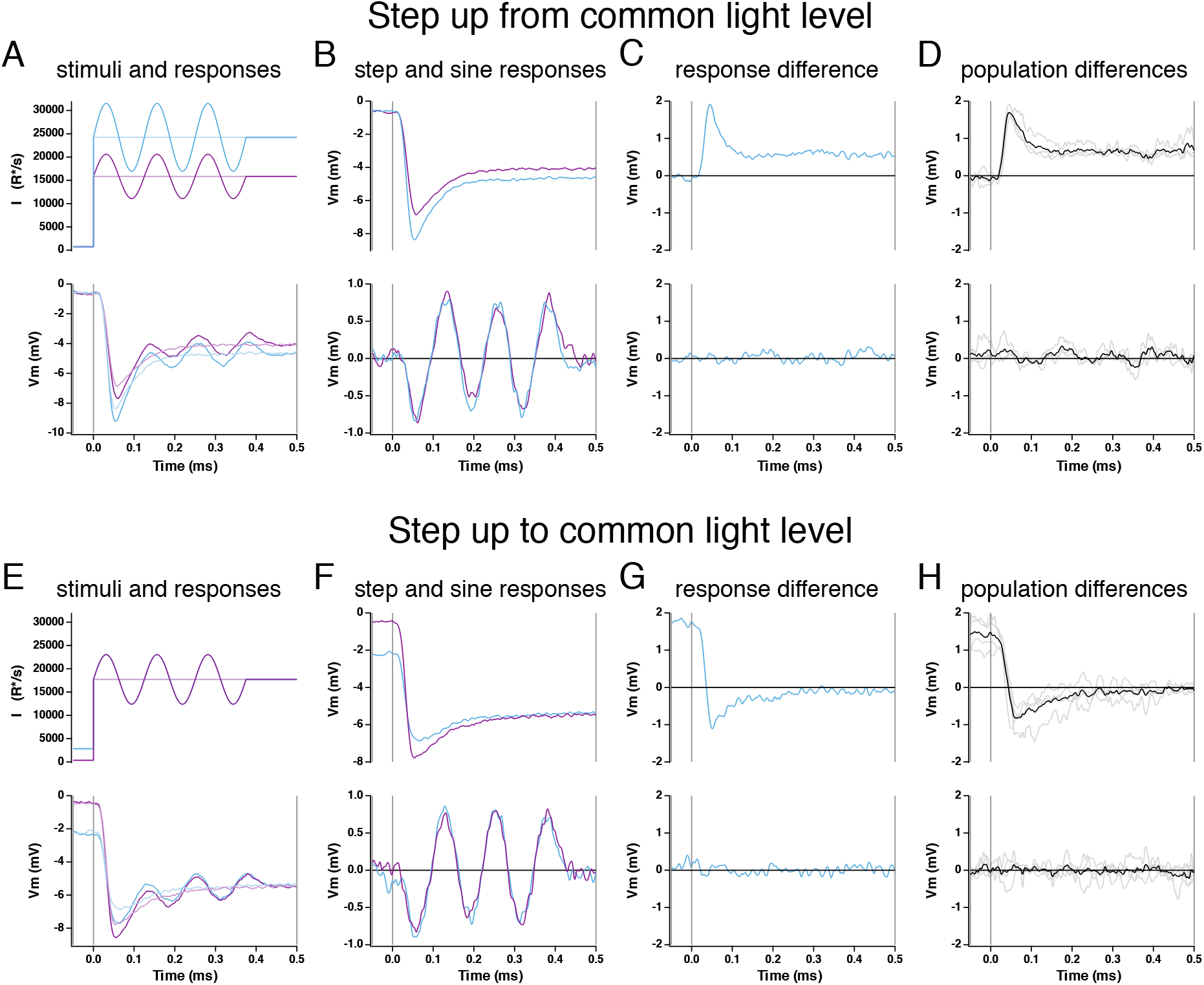
Kinetics of onset of Weber adaptation. **A**. Changes in cone voltage elicited by steps from a common low light intensity to two different high intensities. Superimposed sinusoids probed gain over time following the step in mean intensity. **B**. Voltage responses to the light step alone (top) and to the sinusoidal stimulus (with step response subtracted, bottom). **C**. Difference in step (top) and sine (bottom) responses at the two light intensities. **D**. Mean (± SEM) differences for four cones. **E-H**. As in A-D, but for a light step from two different starting intensities to a common final intensity.

Figure 3E-H show a complementary experiment in which steps were made from two low light levels to a single common high light level. Low light levels were chosen to be below the range where Weber adaptation operates (Angueyra & Rieke, 2013), so that sinusoidal responses at the low light levels would differ more than 2-fold (see Figure 4B). The response to the light step exhibited a dependence on the initial light level, but the sinusoidal response at the high light level did not. Responses that obey Weber’s law were achieved quickly (< 60 ms), well before the response to the step itself had stopped changing and lost its history dependence (∼250 ms). Thus, the onset of Weber adaptation is rapid compared to the step response and to the typical duration of a fixation between saccades; this is consistent with the rapid adaptation observed in Figure 2 for steps and flashes.

**Figure 4.**
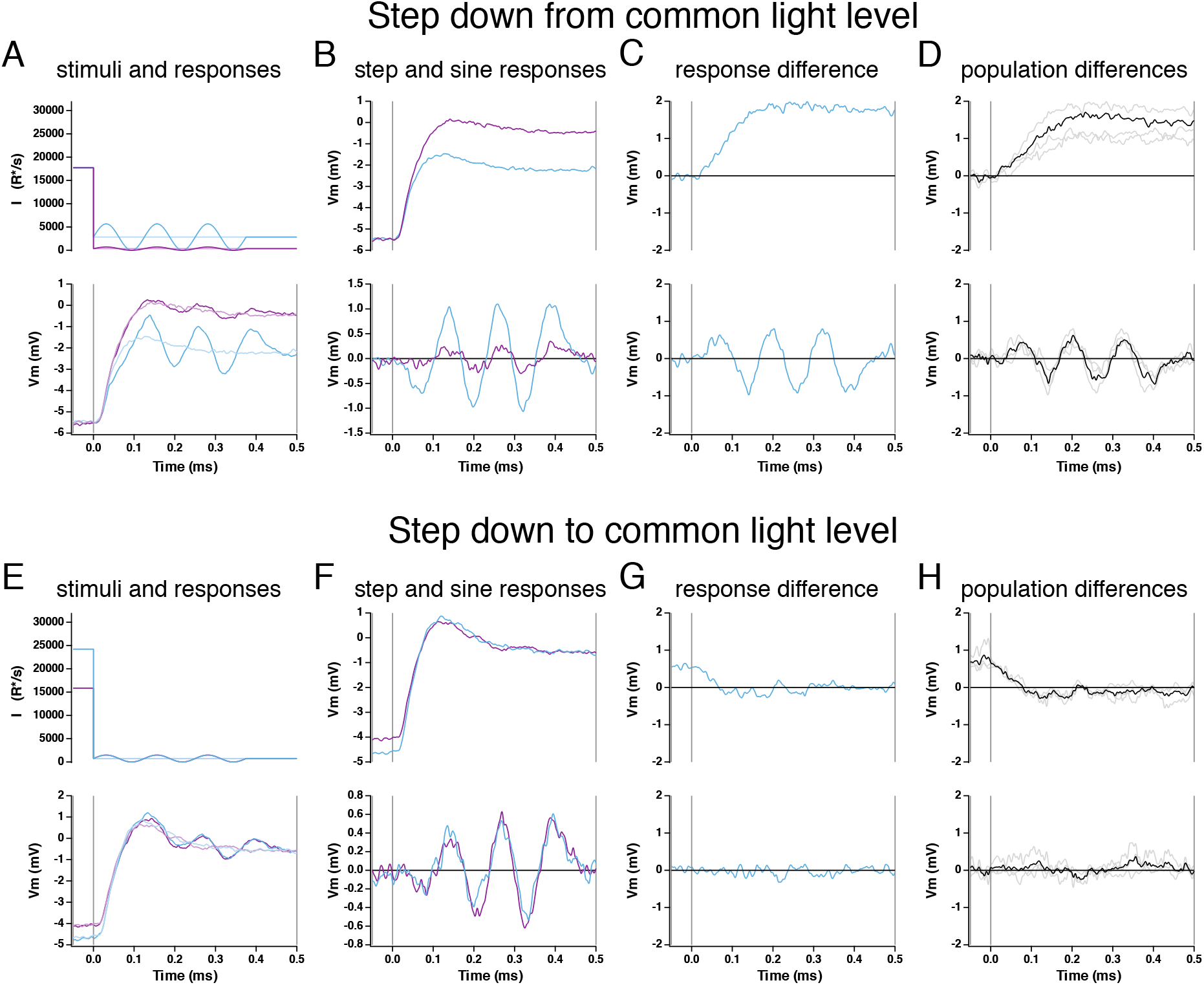
Kinetics of offset of Weber adaptation. **A**. Changes in cone voltage elicited by steps from a common high light intensity to two different low intensities. Superimposed sinusoids probed gain over time following the step in mean intensity. **B**. Voltage responses to the light step alone (top) and to the sinusoidal stimulus (with step response subtracted, bottom). **C**. Difference in step (top) and sine (bottom) responses at the two light intensities. **D**. Mean (± SEM) differences for four cones. **E-H**. As in A-D, but for a light step from two different starting intensities to a common final intensity.

We used a similar approach to probe the kinetics of the offset of Weber adaptation — now measuring responses to steps to different low light levels from a common high light level (Figure 4A) or to a common low light level from different high light levels (Figure 4E). As above, low light levels were chosen to be below the range in which Weber adaptation generates contrast invariance, so that the offset of adaptation caused responses to equal contrast to depend on the mean light level (Angueyra & Rieke, 2013). Isolated responses to the sinusoidal stimuli differed almost immediately following a decrease in mean light level (Figure 4C and D), indicating rapid adaptation to the new mean light level. Similarly, sinusoidal responses measured at a common low light level rapidly lost any dependence on the initial high light level (Figure 4G and H). The sinusoidal responses grew in amplitude for 100-200 ms following the decrease in mean light level, consistent with the kinetics of the recovery of gain for the steps and flashes protocol (Figure 4B and F; compare to Figure 2E and F). The history dependence of the step responses similarly persisted for 100-200 ms.

Figures 2-4 show that the onset of adaptation in responses to both flashes and sinusoidal stimuli is more rapid than the offset and that both are completed within the ∼300-500 ms duration of a single fixation. Further, the time course of the gain changes does not always follow that of the response to the change in mean light level, particularly following increases in light intensity.

#### Responses to light increments and decrements are asymmetric

The asymmetry in the kinetics of adaptation following increases and decreases in mean light level suggested that responses to light increments and decrements might also be asymmetric, as observed in amphibian and fish cones (Baylor & Hodgkin, 1974; Endeman & Kamermans, 2010). This is an important issue because increment/ decrement asymmetries observed in downstream cells are often attributed to differential processing in ON and OFF circuits rather than asymmetric cone signals (see Discussion).

To test for increment/decrement asymmetries in primate cones, we delivered positive and negative steps of equal contrast relative to the background intensity while recording cone photocurrent or photovoltage (Figure 5A and B). Responses to steps with a contrast < 25% were near-symmetric, but responses to higher contrast decrements exceeded responses to increments. This increment/decrement asymmetry was also apparent in the cone voltage responses and the cone synaptic output as measured in recordings from horizontal cells (Figure 5 - Figure Supplement 1). We quantified the increment/decrement asymmetry from the ratio of the mean currents at the end of the 100% contrast steps. The ratio of decrement to increment responses exceeded one across all light levels probed (Figure 5B). The asymmetry was stronger with increasing background intensity (compare Figure 5A top and bottom; p < 0.001 for ratio of asymmetries for step responses from intensities < 6000 R*/s and > 6000 R*/s).

**Figure 5.**
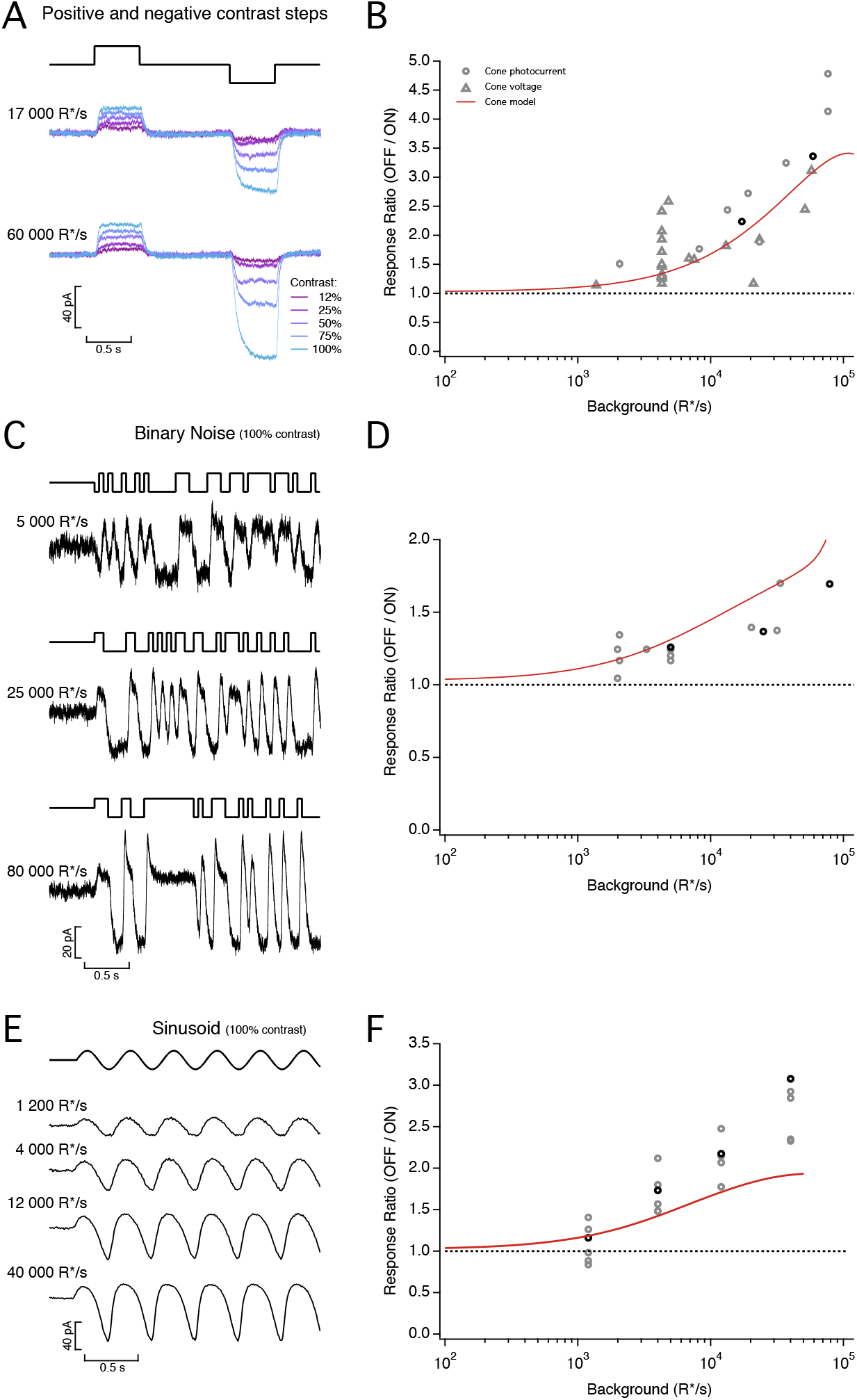
Asymmetric responses to light increments and decrements. **A**. Average cone photocurrents elicited by light increments and decrements at background light intensities of 17,000 R*/s (top) and 60,000 R*/s (bottom). Decrements produced larger responses than symmetric increments for Weber contrasts above 25% (10 traces averaged at each contrast). The asymmetry was larger as background light intensity increased. **B**. Ratio of mean negative to mean positive response to 100% contrast steps as a function of mean light intensity. Red line shows prediction of biophysical cone model. **C**. Photocurrents elicited by a binary noise stimulus (100% contrast) at three mean light intensities. **D**. Ratio of mean negative to mean positive response to binary noise as a function of mean light intensity. Red line shows prediction of biophysical cone model. **E**. Photocurrents elicited by sinusoidal stimuli (temporal frequency 2.5 Hz, 100% contrast) at 4 mean light intensities. **F**. Ratio of peak negative to peak positive response as a function of mean light intensity. Red line is prediction from biophysical cone model.

As an additional test of increment/decrement asymmetries, we stimulated cones with high-contrast binary noise while recording photocurrents. As expected, these stimuli also elicited asymmetric responses, with larger current changes upon decreases in light (Figure 5C). We again quantified the asymmetry as the ratio of mean currents elicited by decrement to increment stimulation; this analysis further confirmed that the asymmetry is stronger as background intensity increases (Figure 5D). Finally, the asymmetry between increment and decrement responses is also clear in responses to high-contrast sinusoidal stimuli (Figure 5E); as for steps, the asymmetry in responses to sinusoidal stimuli increased systematically with increasing mean light level (Figure 5F; p < 1e-4 for sinusoidal stimuli for intensities < 6000 R*/s and > 6000 R*/s).

### A biophysical model of cone responses

The results described above show that primate cones, not unlike cones from other species, have complex responses that cannot be predicted easily from common empirical models. The complexity of these responses originates at least in part from adaptational mechanisms that quickly and strongly adjust cone responses to the prevailing inputs. Below, we test the ability of a biophysical model of cone phototransduction to account for the cone responses illustrated in Figures 1 - 5. In addition to testing the completeness of current understanding of cone phototransduction, our goal was to develop a model that permitted prediction and manipulation of cone responses to a wide range of stimuli.

Two types of models have been used to capture photoreceptor responses. *Empirical models* aim to succinctly capture the dynamics of phototransduction without a tight correspondence with the underlying mechanisms (Clark, Benichou, Meister, & Azeredo da Silveira, 2013; De Palo et al., 2013). Rapid adaptation emerges in these models from feedback or feedforward mechanisms. *Biophysical models* are based directly on the biochemical reactions that comprise the phototransduction process (Younger, McCarthy, & Owen, 1996; Nikonov, Lamb, & Pugh, 2000; Rieke & Baylor, 1998; Korenbrot, 2012; Endeman & Kamermans, 2010; van Hateren, 2005). Rapid adaptation in these models emerges from changes in the rate of cGMP turnover produced by light-dependent changes in phosphodiesterase activity and by calcium feedback to the rate of cGMP production (van Hateren, 2005; Nikonov et al., 2000). We focus here on biophysical models as they captured cone responses at least as well as empirical models.

We represent the enzymatic reactions of the phototransduction cascade as a set of six differential equations. We implement slow adaptation with a slow calcium-dependent feedback that regulates the activity of cGMP channels (Korenbrot, 2012; Rebrik, Botchkina, Arshavsky, Craft, & Korenbrot, 2012; Korenbrot, Mehta, Tserentsoodol, Postlethwait, & Rebrik, 2013) (Figure 6A; see Discussion), but this mechanistic instantiation is not unique. This model has a total of 15 parameters corresponding to time constants, affinities, cooperativities and concentrations of the different components of the phototransduction cascade (see Methods and (Rieke & Baylor, 1998)). Three of these parameters could be expressed in terms of others using steady-state conditions. Six other parameters were measured directly or fixed based on published values, leaving a model with six free parameters that we fit numerically to measured responses to a variety of stimuli (Table 1; see Methods for details).

**Table 1.**
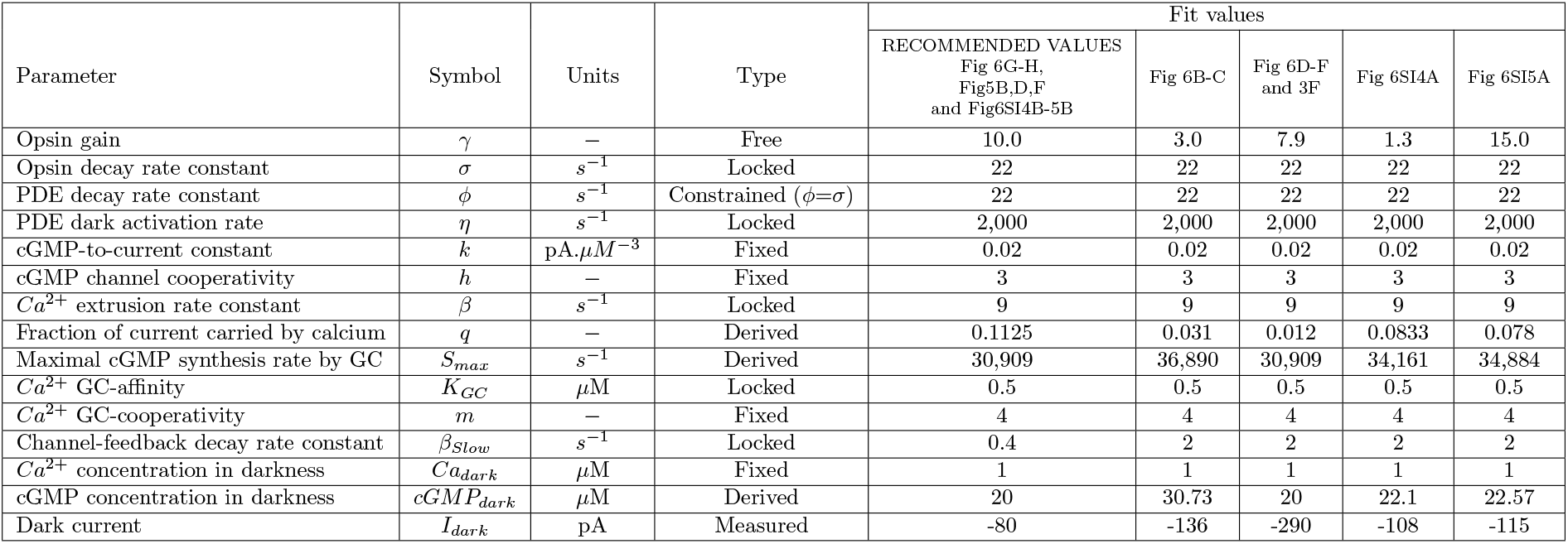
Parameters and best fit values for cone-phototransduction biophysical model.

| Parameter | Symbol | Units | Type | Fit values |  |  |  |  |
| --- | --- | --- | --- | --- | --- | --- | --- | --- |
|  |  |  |  | RECOMMENDED VALUES<br>Fig 6G-H,<br>Fig5B,D,F<br>and Fig6SI4B-5B | Fig 6B-C | Fig 6D-F<br>and 3F | Fig 6SI4A | Fig 6SI5A |
| Opsin gain | $\gamma$ | — | Free | 10.0 | 3.0 | 7.9 | 1.3 | 15.0 |
| Opsin decay rate constant | $\sigma$ | $s^{-1}$ | Locked | 22 | 22 | 22 | 22 | 22 |
| PDE decay rate constant | $\phi$ | $s^{-1}$ | Constrained ($\phi=\sigma$) | 22 | 22 | 22 | 22 | 22 |
| PDE dark activation rate | $\eta$ | $s^{-1}$ | Locked | 2,000 | 2,000 | 2,000 | 2,000 | 2,000 |
| cGMP-to-current constant | $k$ | $pA \cdot \mu M^{-3}$ | Fixed | 0.02 | 0.02 | 0.02 | 0.02 | 0.02 |
| cGMP channel cooperativity | $h$ | — | Fixed | 3 | 3 | 3 | 3 | 3 |
| $Ca^{2+}$ extrusion rate constant | $\beta$ | $s^{-1}$ | Locked | 9 | 9 | 9 | 9 | 9 |
| Fraction of current carried by calcium | $q$ | — | Derived | 0.1125 | 0.031 | 0.012 | 0.0833 | 0.078 |
| Maximal cGMP synthesis rate by GC | $S_{max}$ | $s^{-1}$ | Derived | 30,909 | 36,890 | 30,909 | 34,161 | 34,884 |
| $Ca^{2+}$ GC-affinity | $K_{GC}$ | $\mu M$ | Locked | 0.5 | 0.5 | 0.5 | 0.5 | 0.5 |
| $Ca^{2+}$ GC-cooperativity | $m$ | — | Fixed | 4 | 4 | 4 | 4 | 4 |
| Channel-feedback decay rate constant | $\beta_{Slow}$ | $s^{-1}$ | Locked | 0.4 | 2 | 2 | 2 | 2 |
| $Ca^{2+}$ concentration in darkness | $Ca_{dark}$ | $\mu M$ | Fixed | 1 | 1 | 1 | 1 | 1 |
| cGMP concentration in darkness | $cGMP_{dark}$ | $\mu M$ | Derived | 20 | 30.73 | 20 | 22.1 | 22.57 |
| Dark current | $I_{dark}$ | pA | Measured | -80 | -136 | -290 | -108 | -115 |

**Figure 6.**
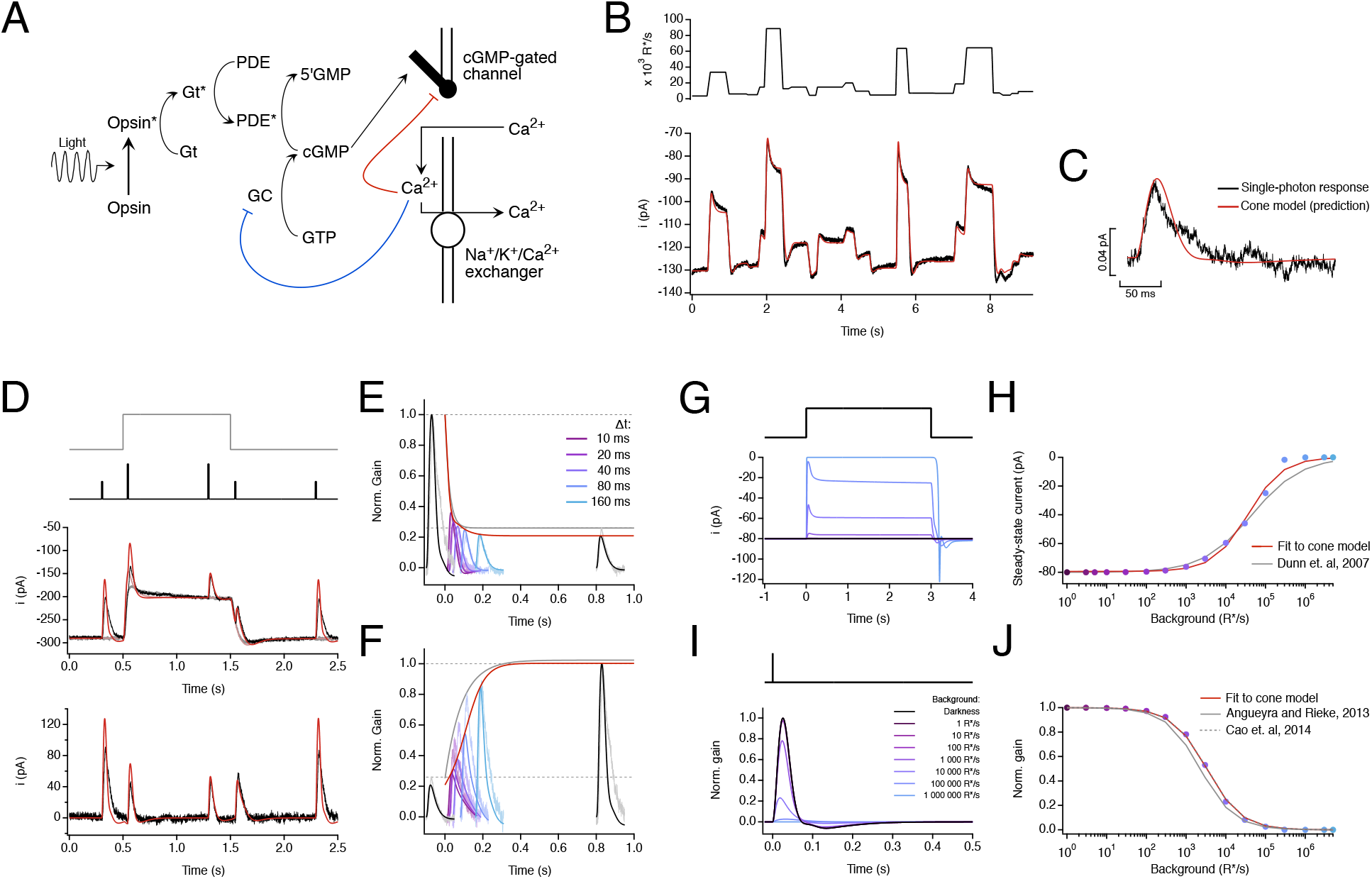
A biophysical model of phototransduction captures a wide range of the cone responses. **A**. Schematic of phototransduction cascade and corresponding components of the biophysical model: cyclic-GMP (cGMP) is constantly synthesized by guanylate cyclase (GC), opening cGMP-gated channels in the membrane. Light-activated opsin (Opsin*) leads to channel closure by activating the G-protein transducin (Gt*) which activates phosphodiesterase (PDE*) and decreases the cGMP concentration. Calcium ions (Ca^2+^) flow into the cone outer segment through the cGMP-gated channels and are extruded through Na^+^/K^+^/Ca^2+^ exchangers in the membrane. Two distinct feedback mechanisms were implemented as calcium-dependent processes that affect the rate of cGMP-synthesis (blue line) and the activity of the cGMP-gated channels (red line). **B**. Fit to the measured cone response to the naturalistic stimulus shown in Figure 1. The model is able to capture both the currents at the end of fixations and the response transients following rapid changes in the stimulus. See Table 1 for fit parameters. **C**. The model accurately predicts the amplitude and kinetics of the single-photon response. **D-F**. Model fit to step and flashes responses from Figure 2. The model exhibits fast changes in gain at step onset (τ_On-Model_ = 13.6 ms) and a slower recovery of gain at step offset (τ_Off-Model_ = 180 ms). These time constants are compared to experimental data in Figure 2F. **G**. Model responses to steps of increasing intensity. **H**. Dependence of model’s steady-state current on background light intensity (colored dots). This relation was fit with a Hill equation (Dunn *et al*., 2007) with a half-maximal background, I_½_ = 43,500 R*/s and a Hill exponent, n = 0.77. The fit obtained by Dunn *et. al*., 2007 (I_½_ = 45,000 R*/s and n = 0.7) has been replicated for comparison (gray line). **I**. Estimated single-photon responses of the model, normalized by the response in darkness, at increasing background light intensities. **J**. Relation of the model’s peak sensitivity, normalized to the peak sensitivity in darkness, across background light-intensity (colored dots). The half-desensitizing background (I_0_) for the model is 3297 R*/s. The fits obtained in Angueyra and Rieke, 2013 (I_0_ = 2250 R*/s, after correcting a calibration error in the original article) and in Cao *et al*., 2014, (I_0_ = 3330 R*/s, assuming a collecting area of 0.37 μm^2^ for transversally illuminated cones) have been replicated for comparison (gray lines).

Because of the limited duration of our recordings, we could not measure all the responses used in fitting from the same cone. Using responses from several cones simultaneously in model fitting required accounting for differences in sensitivity and dark current between cones. Dark current and sensitivity are set by the dark cGMP concentration and the gain of photopigment activation (the ‘opsin gain’). Hence, we tested the ability of the model to generalize across cones and stimuli by allowing these two parameters to vary, while keeping the remaining parameters fixed. This procedure means that the parameters determining the kinetics of the model responses are consistent across all fitted cones. Several approaches to fitting model parameters provided very similar results (see Methods for details).

Figure 6B compares measured and model responses to the naturalistic stimulus from Figure 1. The model successfully captures the dynamic changes in current and the final current at the end of each fixation (Figure 6B). The contribution of slow adaptation was relatively minor, but it did help capture responses to long light steps, including the fixations in Figure 1 (compare to Figure 6 - Figure Supplement 1). Empirical models with a comparable number of free parameters could also capture responses of individual cones to naturalistic stimuli (Figure 6 - Figure Supplement 2-3).

The model was also able to fit responses to other stimuli after adjustments to dark current and sensitivity to account for differences between cones. First, the model correctly predicted the amplitude and kinetics of the single-photon response (Figure 6C). Second, the model captured both the slow dynamics of responses to light steps and the fast changes in the amplitude and kinetics of flash responses superimposed on these steps (Figure 6D). As for real cones, the offset of adaptation in the model was slower than the onset (Figure 6E, and red triangle in Figure 2F). Third, the model captured changes in cone steady-state current and sensitivity across a wide range of light levels, with model responses showing Weber adaptation close to that measured (Figure 6G-H) (Dunn et al., 2007; Angueyra & Rieke, 2013; Cao et al., 2014). Fourth, the model exhibited asymmetric responses to light increments and decrements that fall within the range of the data (Figure 5B, D and F, red lines). Empirical models did not generalize as well; in particular, these models struggled to capture the changes in steady-state current and the dependence of response gain on background (Figure 6 - Figure Supplement 2-5; see Discussion). These models lack an intrinsic baseline or “dark” activity that controls the onset of adaptation. Hence adaptation operates at all light levels, and this causes a failure to generalize to responses to stimuli for which adaptation contributes little - such as flashes delivered in darkness.

The biophysical model described above, while not perfect, captures cone responses to a broad range of stimuli. The success of the model indicates that the known operation of cone phototransduction can explain cone responses to the highly dynamic inputs encountered during natural vision. The model allows us (1) to predict how signals in the cone array encode a variety of inputs and (2) to manipulate cone responses, e.g. to remove the effects of adaptation. Below we provide examples of each of these applications of the model.

### Applications of biophysical model to neural coding

#### Local vs global adaptation

Most existing models for ganglion-cell responses share a common architecture in which retinal inputs are first processed linearly over space and time, followed by a nonlinear processing step associated with bipolar synapses or spike generation in ganglion cells (Ozuysal & Baccus, 2012; Pillow et al., 2008; Cui, Wang, Park, Demb, & Butts, 2016). For these models to be effective, they must either be restricted to stimuli for which the cones do not adapt, or adaptation in the cones must be accounted for by the late nonlinear steps in the model. But adaptation operates independently within each cone and hence is spatially local, unlike post-cone circuit mechanisms that likely have access only to signals pooled across multiple cones due to convergence of cone signals in retinal circuits. The cone model described above provides an opportunity to identify visual inputs for which the spatial locality of adaptation may have an important role in shaping retinal signals.

To identify such input stimuli, we compared a model in which adaptation occurred prior to pooling of signals across cones (‘*cone-adaptation model*’), with a model in which adaptation operated only on the pooled cone signal (‘*post-cone-adaptation model*’, Figure 7A). We presented each model with flashed patches of natural images and compared the predicted responses (Figure 7B). When all the cones encounter similar changes in input (i.e. spatially homogeneous bright or dark image patches such as those encountered in a patch of sky or a tree trunk), the location of adaptation did not matter (Figure 7B, bottom left and middle). This is expected intuitively since in these cases integration of cone signals does not involve averaging heterogeneous cone signals, and hence the pooled cone signal is very similar to the signal present at each cone. Hence adaptation is consistent across cones in the cone-adaptation model, and can be closely replicated by adaptation mechanism in the post-cone-adaptation model. Spatially-structured patches (e.g. patches with tree branches or leaves), however, led to considerable differences in the models with cone and post-cone adaptation (Figure 7B, bottom right). These differences originate because adaptation causes cone responses to equal and opposite light increments and decrements to differ both in steady-state levels and in the kinetics with which they reach that level. Thus, when responses of a cone exposed to a light increment and a cone exposed to an equal and opposite decrement are summed, the steady-state responses partially cancel but the time required for each cone to reach steady state differs due to the different kinetics of adaptation. These effects are created by differences in the inputs to individual cones, and hence cannot be captured by adaptation occurring after integration of cone signals. Capturing such local adaptation will likely be an important aspect of creating predictive models for natural inputs (see Discussion).

**Figure 7:**
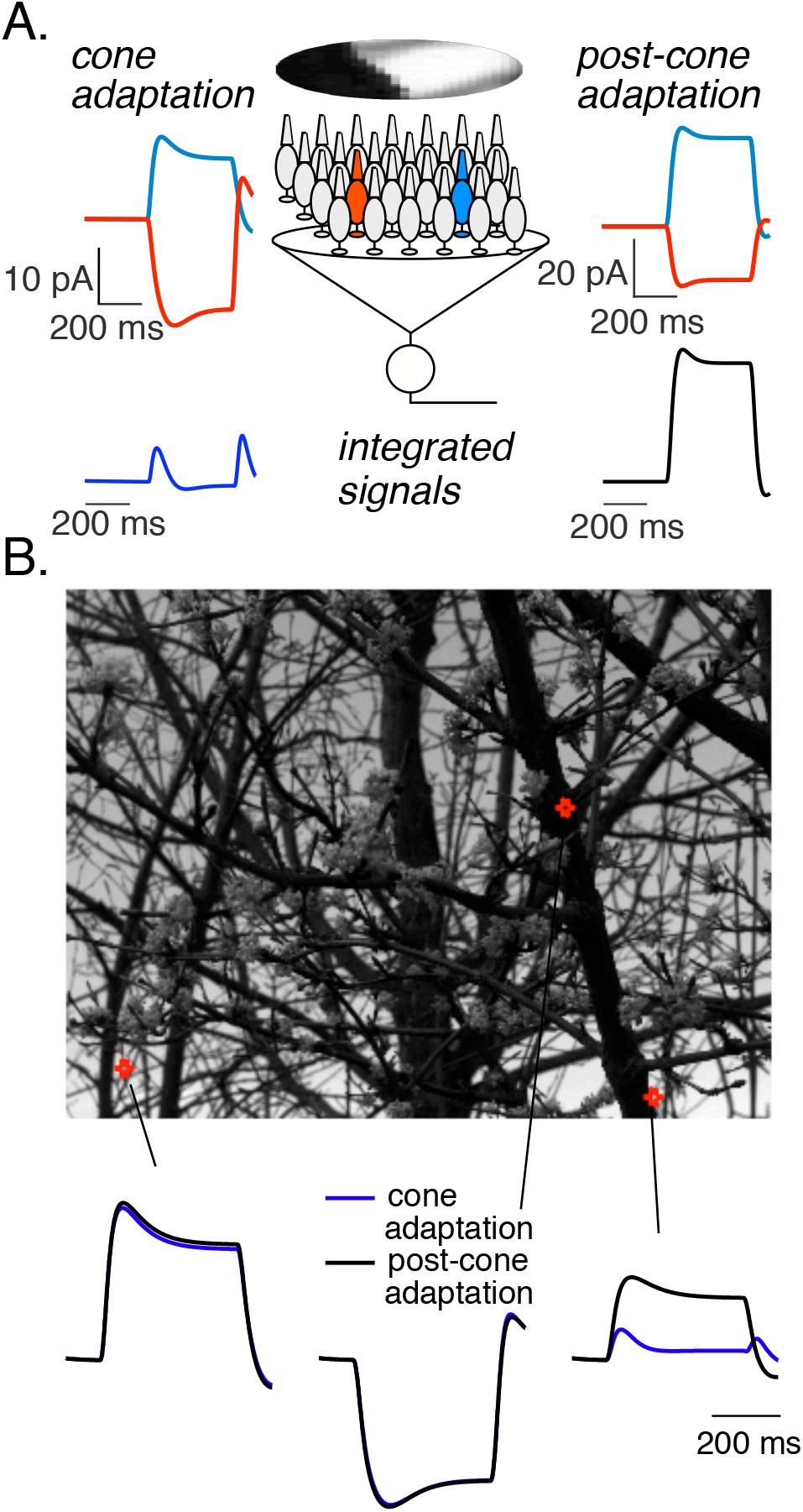
Local cone adaptation shapes integrated responses to spatially structured inputs. A. (top) Examples of predicted responses of two cones to a flashed natural image. Left panels show predicted responses from the biophysical model, while right panels shows predicted responses for a linear cone model. (bottom) Sum of the responses across a collection of cones to illustrate the impact of signal integration - e.g. integration within the receptive field of a downstream neuron. Cones were weighted with a gaussian spatial profile resembling the receptive field of a primate Parasol retinal ganglion cell. The gaussian SD was 10 cone spacings, meaning that receptive field encompassed several hundred cones. Adaptation in the ‘post-cone-adaptation’ model operated on the integrated signal, and responses of individual cones depended linearly on light input. B. Predictions of integrated responses for several image patches. The top panel shows the locations of the illustrated patches. The bottom panel shows the integrated responses for adapting (blue) and non-adapting (black) cones.

#### Manipulating cone responses

In addition to predicting the contribution of cones to responses of downstream neurons, the model described in Figure 6 provides a tool to manipulate specific aspects (e.g. nonlinearities or kinetics) of the cone responses. This will, for example, provide a tool to test the impact of local cone adaptation (as in Figure 7), and more generally to isolate the impact of post-cone circuit nonlinearities on retinal responses.

Figure 8 shows how the cone model can be used to manipulate light stimuli to compensate for adaptation and create linear responses - a procedure we refer to as ‘light-adaptation clamp.’ To accomplish this, we compare the outputs of two cone models: a linear model with the original stimulus as input, and the full model with a transformed version of the original stimulus as input (Figure 8A). The linear cone model is determined by the responses of the full model to a brief low-contrast flash (i.e. the linear range impulse response of the full model; see Methods). The output of the linear model provides the desired response. We then adjust the transformation of the stimulus to minimize the difference between the two models - i.e. to cause the output of the full model to the transformed stimulus to match the output of the linear model to the original stimulus. Figure 8 - Figure Supplement 1 shows an example of several steps in this adjustment process.

**Figure 8:**
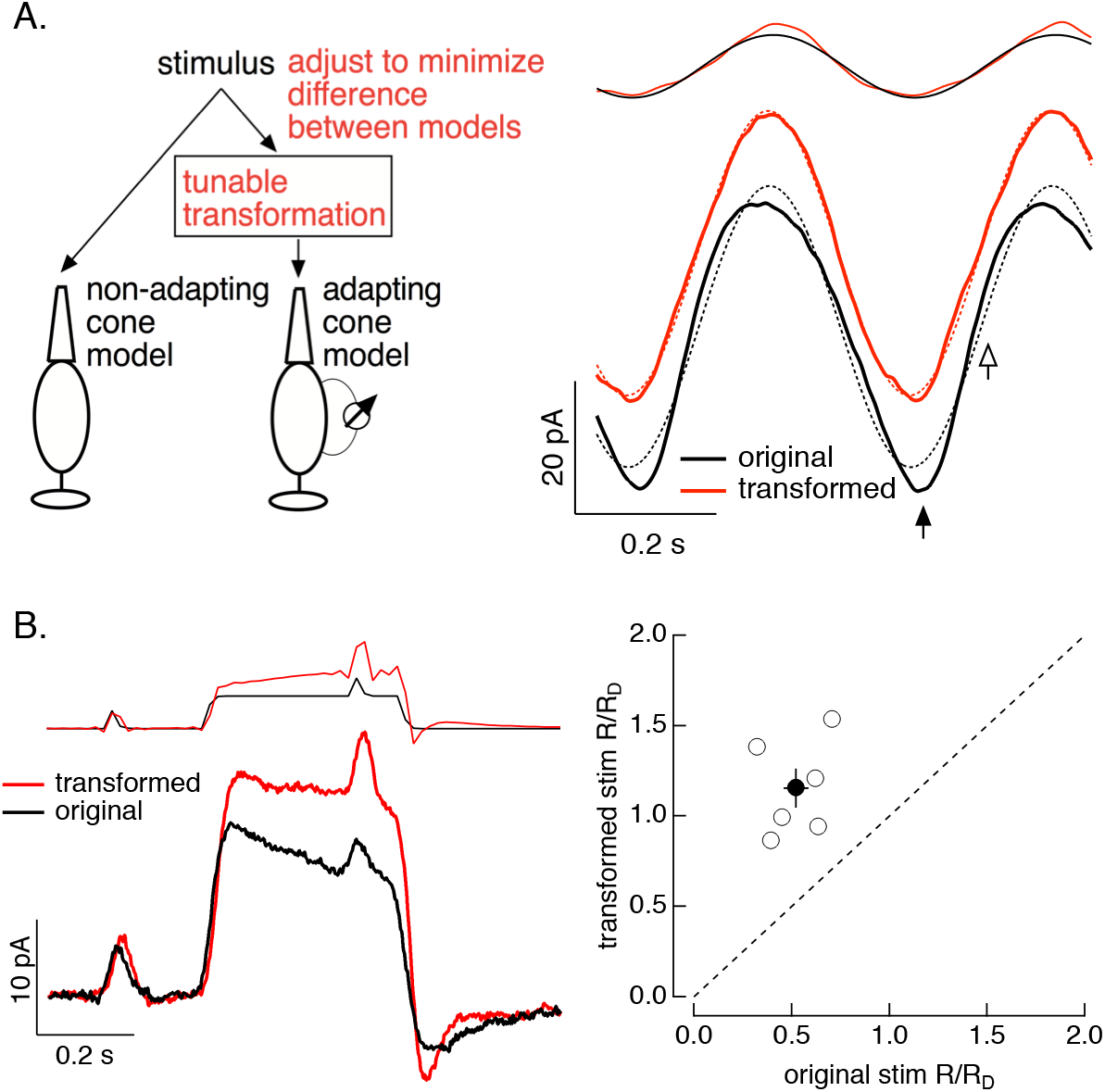
Light-adaptation clamp. **A**. (left) Illustration of procedure. The stimulus to the full cone model is tuned until the output of this model matches the “target” output of a linear (non-adapting) cone model. (right) Example of application to sinusoidal stimuli. The original stimulus and response are shown in black, and the modified stimulus and response in red. Dashed lines show best fit sinusoids. **B**. Application to a step and flash stimulus. Left shows an example cell, right collects data across several cells, plotting the ratio of the amplitude of the responses to flashes before and on top of the step for transformed stimuli (y-axis) and original stimuli (x-axis). The discrete nature of the stimuli originates because these stimuli were delivered using a computer monitor with a 60 Hz frame rate.

The light-adaptation clamp procedure is particularly simple for sinusoidal stimuli. Cone responses to high-contrast sinusoids are far from sinusoidal due to rapid adaptation (see Figure 5E and Figure 8A, right). For sinusoidal stimuli, the output of the linear model is also a sinusoid and hence the light-adaptation clamp procedure identifies a stimulus that makes the output of the full cone model sinusoidal. Measured responses to sinusoidal stimuli differ from a sinusoid in two ways, which are clear when comparing the measured response to a sinusoidal fit (solid and dashed lines in Figure 8A, right): (1) responses to decrements are larger than increments (closed arrow in Figure 8A, right; see also Figure 5E), and (2) the response to the dark-to-light transition is more rapid than expected from a sinusoid (open arrow in Figure 8A, right). These effects are relatively subtle for the light level used in Figure 8, and testing whether we can indeed effectively minimize them is a strong test of the light-adaptation clamp procedure. As expected from the small deviations of the responses from linearity, the predicted transformation to produce linear responses is also subtle (red and black traces at top of Figure 8A, right). Nonetheless, measured cone responses to the transformed stimulus are considerably closer to a sinusoid then responses of the same cones to the original stimulus (black traces and fit in Figure 8A).

This approach is not limited to subtle manipulations of cone signals. Figure 8B tests the ability of the light-adaptation clamp to identify stimuli that minimize adaptation in a steps and flashes protocol similar to the one used in Figure 2 to characterize cone adaptation. As in Figure 2, adaptation considerably reduces the gain of responses to flashes delivered on top of a step compared to those delivered prior to the step. The cone model predicts that a sizable transformation of the original stimulus is needed to minimize this effect of adaptation and obtain the same flash response before and during the step (red and black traces at top of Figure 8b, left). Measured responses to the original and transformed stimuli show that adaptation is indeed largely eliminated by the transformed stimuli, a finding that holds across cones (Figure 8B, right).

It is important to note that this procedure directly tests the model’s ability to generalize across stimuli and across cones, as the stimulus manipulations were predicted from the model fits shown in Figure 6 and subsequently tested in “*naive*” cones. If the model predictions inaccurately captured responses of the measured cones, the procedure should fail to identify manipulations that achieve the desired transformation of the cone signals (linearization in the examples here). The examples above show that we can make predictable manipulations of both subtle and non-subtle aspects of the cone signals and verify that those manipulations indeed work as predicted in cones that did not contribute to the model parameters. These stimulus manipulations provide a tool to manipulate cone signals and establish a causal relationship between their properties and those of responses in downstream neurons (see Discussion).

## Discussion

All sensory systems share a need to adapt to the statistics of the natural environment. Vision is no exception, as we are able to see across ambient light levels differing by more than a factor of one trillion. Visual adaptational mechanisms span a wide range of temporal and spatial scales. Some mechanisms tune circuits and control sensitivity over the course of minutes or hours (e.g. circadian regulation of retinal gap junctions (Bloomfield & Volgyi, 2009; Ribelayga, Cao, & Mangel, 2008) or of opsin expression (von Schantz, Lucas, & Foster, 1999; Li et al., 2005)). More rapid mechanisms operate in less than one second and permit effective encoding of the large and rapid changes in input experienced during free viewing of natural scenes. These mechanisms must balance the need for high sensitivity with the risk of saturation (Abrams, Hillis, & Brainard, 2007; Wark, Fairhall, & Rieke, 2009). The magnitude of this challenge is obvious when trying to take pictures with a digital camera: no single exposure setting can capture the range of inputs encountered in typical visual scenes.

The visual system deals with a significant fraction of these challenges upfront, by imposing fast and local adaptation at every “pixel” or photoreceptor.

### Predictive model for cone signals

Evaluating the contributions of cones to visual function requires developing models that can predict responses to a wide range of stimuli. Functional models for ganglion cell responses often use linear or linear-nonlinear models for cones (Burton, 1973; MacLeod, Williams, & Makous, 1992; Stockman, Petrova, & Henning, 2014; Pillow et al., 2008; Ozuysal & Baccus, 2012); such models do not accurately capture cone responses, particularly the dynamics of responses following large changes in input, such as those occurring following saccades.

Alternatives to linear or linear-nonlinear models include models that incorporate feedback or parallel feedforward signals that can capture history-dependent effects such as adaptation. These models can capture many nonlinearities in cone responses well (Clark et al., 2013). Another approach is to construct mechanistically-based models that reflect the underlying biochemistry of phototransduction. This is the approach we follow here, in part because such models generalized across stimuli better than empirical models (Figure 6), and in part because they permitted a direct test of how well current understanding of phototransduction accounts for responses to a wide range of stimuli.

Several limitations of our model are important to emphasize. First, our model omits several known mechanistic features for simplicity, notably feedback to the opsin and photopigment bleaching. These mechanisms are important at higher light levels and have been included in other biophysical models (Lamb & Pugh, 2004; van Hateren & Snippe, 2007). Second, we chose a set of model parameters that provided a good fit to our measured responses. However, some parameters in the model can trade against each other, meaning that more than one combination of parameters can provide a good fit. As a consequence, model parameters should not be interpreted as unique estimates of actual biochemical rate constants.

The biophysical model presented here captures cone responses to a broad range of stimuli within the range of mean intensities we tested (up to ∼100,000 R*/s for short periods of time). This spans the range of light levels explored in most physiological and psychophysical studies of cone vision. As a result, the model can be used to explore the impact of cone signaling on downstream visual responses, as described below.

### Separation of cone and post-cone processing to visual function

The model we develop provides a needed tool to evaluate the contributions of the cones and post-cone processing to shaping of responses in subsequent visual neurons in the retina or cortex. Asymmetric sensitivities to contrast increments and decrements provide one example. These asymmetries are a common feature of responses of retinal ganglion cells and V1 cortical neurons and of behavior. Such asymmetries are often attributed to the circuits that read out the cone responses, with the implicit assumption that the cones provide symmetrical input to ON and OFF circuits. In some cases this is almost certainly accurate. For example, psychophysical thresholds for detecting rapid contrast decrements (stimuli designed to isolate the OFF pathway) are lower than those for detecting rapid contrast increments (designed to isolate the ON pathway) (Bowen, Pokorny, & Smith, 1989). Because detection of these stimuli requires just a few percent contrast (where cone responses are near linear), the difference in sensitivity almost certainly arises in the post-cone circuitry. At higher contrasts, decrements elicit larger V1 responses than increments in humans and monkeys (Kremkow et al., 2014), and detection of 100% contrast decrements embedded in binary noise is more reliable than detection of 100% contrast increments (Komban et al., 2014). These high-contrast stimuli will elicit asymmetric responses in the cones themselves, and cone nonlinearities likely contribute substantially to downstream signaling.

The cone light-adaptation clamp procedure we introduce here could help reveal the contribution of the cones to these (and other) downstream signals. As illustrated in Figure 8, this approach permits identification of stimuli that generate desired cone responses — e.g. symmetrical responses to increments and decrements. The use of such stimuli while recording responses of downstream visual neurons or while monitoring perception should help separate the contributions of cones from those of post-cone circuits. Indeed, we have used the light-adaptation clamp approach to show an unexpected role of cone adaptation in how some ganglion cell types respond to spatial structure in natural inputs (Z. Yu, M. Turner, F. Rieke, unpublished).

### Importance of cone adaptation for models of signaling in retinal ganglion cells

The past 20 years has seen a dramatic advance in our understanding of what information retinal ganglion cells provide to central targets. As a result, ganglion cells serve as a leading example of how connectivity and signaling mechanisms shape the outputs of a neural circuit in a behaviorally-important manner (reviewed by (Field & Chichilnisky, 2007; Sanes & Masland, 2015)). Nonetheless, current models for ganglion-cell feature selectivity generalize poorly to novel stimuli, particularly naturalistic ones (Heitman et al., 2016; McIntosh, Maheswaranathan, Nayebi, Ganguli, & Baccus, 2016). This failure to generalize may occur at least in part because current models lack adaptation in individual cones, instead assuming that the cones respond linearly across stimuli. Yet, as we show here, naturalistic stimuli strongly engage adaptation in the cones (Figure 1). These considerations suggest that cone adaptation, and its natural operation on a small spatial scale, will be a key factor shaping retinal output signals. For example, our cone model predicts that patches of natural images with high spatial structure will produce transient responses when signals from multiple cones are integrated (Figure 7).

Better models for ganglion-cell function are needed for improved retinal prosthetics and to determine how key steps in visual processing are distributed between retinal and cortical mechanisms. A potential limitation of current models is that they do not reflect the functional architecture of the underlying circuits - particularly with respect to the location of key circuit nonlinearities. Local adaptation is just one example in which getting the order of linear and nonlinear steps correct matters for predicting output responses. Incorporating the model that we develop here for nonlinear, adaptive cone signaling into models for downstream visual neurons could be an important step towards models that generalize across stimuli.

## Methods

### Animals, tissue and solutions

We made electrophysiological recordings from primate retinas (*Macaca fascicularis, nemestrina* and *mulatta* of either sex, ages 3 – 19 yrs) in accordance with the Institutional Animal Care and Use Committee at the University of Washington. We obtained retina through the Tissue Distribution Program of the Regional Primate Research Center. Most enucleations were performed under pentobarbital anesthesia; a few were performed with halothane anesthesia. After enucleation, we rapidly (< 3 min) separated the retina-pigment epithelium-sclera complex from the anterior segment, removed the vitreous humor, and dark-adapted the retina for 1h in warm (32° C) Ames medium bubbled with a mixture of 95% CO_2_ and 5% O_2_. In some young animals, we removed the vitreous after incubation in plasmin (∼50 μg/mL in ∼10mL of solution for ∼20 min at room temperature). We performed all subsequent procedures under infrared illumination (> 900 nm). For recording, we separated a small piece of retina (∼4 mm^2^) from the pigment epithelium and mounted it, photoreceptor-side up, on a poly-lysine coated coverslip (BD Biosciences) that formed the floor of a recording chamber. We continually superfused retinas with warm (∼31°-33° C) oxygenated Ames medium. Treatment with DNase I (Sigma-Aldrich) (30 units in ∼250 μL of Ames for 4 min) facilitated access to the photoreceptor inner segments. For horizontal-cell recordings, we obtained thin vibratome slices (∼200 μm) using chilled Ames-medium. Subsequently, individual slices were transferred to warm bicarbonate-buffered Ames-medium for storage until recording. All the recordings presented here were made in peripheral retina (>20° eccentricity).

### Recordings, light-stimulation and analysis

For whole-cell voltage-clamp recordings of cone photoreceptors (junction-corrected holding potential –70 mV) we measured signals with an internal solution containing (in mM): 133 potassium aspartate, 10 KCl, 10 HEPES, 1 MgCl_2_, 4 Mg-ATP, 0.5 Tris-GTP; pH was adjusted to 7.2 with NMG-OH and osmolarity was 280 ± 2 mOSM. The internal solution did not contain any calcium buffer (or calcium), as even low concentrations of calcium buffer caused the cone light response to become increasingly biphasic during the course of a recording. For perforated-patch current-clamp recordings (without current-injection), we used an internal solution containing (in mM): 125 potassium aspartate, 10 KCl, 10 HEPES, 5 EGTA, 1 MgCl_2_, 0.5 CaCl_2_, 4 Mg-ATP, 0.5 Tris-GTP, and 30 µg/mL gramicidin; pH was adjusted to 7.2 with NMG-OH and osmolarity was 280 ± 2 mOSM. We included EGTA in the internal so that inadvertent whole-cell access caused responses to rapidly become biphasic; any such recordings were terminated. For whole-cell recordings from horizontal cells we used the same internal solution used for the perforated-patch recordings, but omitted gramicidin. Access resistance was ∼10-14 MΩ and was compensated 50% (prediction and compensation settings on Multiclamp 700B Amplifier).

Light stimuli from blue, green and red LEDs (peak wavelengths 405, 510 and 640 nm) permitted quick identification of cone types. The stimuli illuminated a ∼150 μm diameter area centered and focused on the recorded cone through the condenser of an upright microscope. We converted photon densities (photons/μm^2^) to R*/photoreceptor using a collecting area of 0.6 μm^2^ (Schneeweis & Schnapf, 1999), previously measured cone spectral sensitivities (Baylor, Nunn, & Schnapf, 1984) and measured LED spectra. For horizontal-cell recordings, the stimuli illuminated a ∼500 μm diameter area, and we assumed a collecting area of 0.37 μm^2^ (Schnapf et al., 1990), as illumination was incident directly on the outer segments instead of funneled through the inner segments. We calibrated stimuli using M-cones as reference. The M-cone:L-cone:S-cone:rod sensitivity ratio for the blue LED used during the horizontal-cell recordings is 1:0.48:1.37:3.25. Horizontal cells were adapted for at least 2 minutes to a background 350 R*/M-cone/s before data collection (equivalent to ∼1100 R*/rod/s or ∼175 R*/L-Cone/s).

We acquired data using Multiclamp 700B amplifiers. We low-pass filtered recorded currents at 3 kHz and digitized the data at 20 kHz. After analysis, we digitally low-pass filtered the data in Figures 3 and 4 at 200 Hz for ease of viewing. We analyzed recorded data through custom routines in Matlab (The Mathworks). We excluded data from cells that showed unusually rapid run-down of light responses, low sensitivity or from short-lived recordings. All cones presented here were either L or M cones.

### Model of fixation duration and saccades

The model to generate naturalistic stimuli was based on a statistical approximation from measurements of eye movements made in humans (Harris et al., 1988). Fixation times in this model follow an exponential distribution with a refractory period, such that the probability of two saccades following each other is given by:

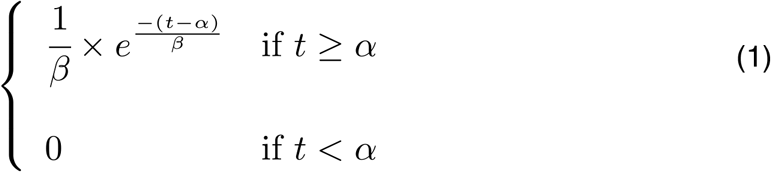

with α, the refractory period, of 100 ms (Harris et al., 1988); a value of β, the time constant of the exponential distribution, of 200 ms was selected to generate on average 3 saccades every second (range of 2 to 5 saccades/s). This model did not include any fixational eye movements. Saccades were inserted between fixations, with a duration (D_S_), proportional to a random amplitude (A_s_), and dictated by:

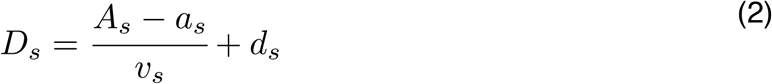

where the velocity, v_s_, was drawn from a uniform distribution between 0.4 and 0.6°/ms, a_s_ = 10° and d_s_ = 40 ms (Rucci, Edelman, & Wray, 2000). This generated saccades that lasted ∼15 ms. At each fixation the intensity was drawn from distributions constructed from individual natural images taken from the van Hateren database (van Hateren & Snippe, 2007). During saccades, transitions between intensities were linear.

### Linear and linear-nonlinear model

Linear filters corresponded to estimates of the single-photon response, obtained by recording cone responses to dim flashes in darkness and dividing them by the strength of the flash. For dim flashes in darkness we chose flash intensities between 100 and 200 R*/flash; the flash intensity for the cell in Figure 1 was 177 R*/flash. The single-photon responses were then fitted with the following equation (Angueyra & Rieke, 2013; Baylor, Nunn, & Schnapf, 1987):

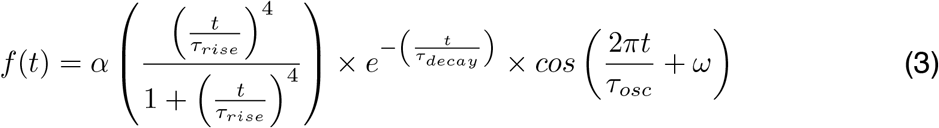

The parameters for the example cell in Figure 1c were: α = 631, τ_rise_= 28.1 ms, τ_decay_= 24.3 ms, τ_osc_ = 2×10^3^ s and ω = 89.97°. This cell did not show a significant oscillation in its response and hence the cosine term in the fit was not needed; others cells did show small undershoots that were better fit when we included the cosine term (see also (Angueyra & Rieke, 2013)).

The linear filter was convolved directly with the light stimulus to obtain a linear estimate of the responses. Given that the linear filter was obtained in darkness, where gain is maximal, we allowed rescaling of the linear model by a single factor. The rescaling factor was chosen to match the current at the end of the highest fixation, and had a value of 0.01 for the example cone presented in Figure 1.

After baseline subtraction, the relationship between the real and linear model currents (mean current during the final 50 ms of each fixation) was fitted with the following function:

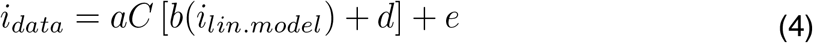

where C[] is the cumulative density of a normal function with zero mean and standard deviation of 1 (Chichilnisky, 2001). The parameters for the fit shown in Figure 1D were a = 305.4, b = 0.039, d = 1.00 and e = −262.9.

### Biophysical model of cone phototransduction

The biophysical model of the phototransduction cascade briefly presented here (Figure 6) is a modification of a model of phototransduction for toad rods (Rieke & Baylor, 1996; Rieke & Baylor, 1998). The original rod model is largely equivalent to other biophysical models successfully used in the past in rods and cones from other species (Pugh & Lamb, 1993; Nikonov, Engheta, & Pugh, 1998; Endeman & Kamermans, 2010), and as the first component of a primate horizontal-cell model (van Hateren, 2005). In the model, adaptation emerges through activity-dependent changes in the cGMP turnover, produced by a light-induced increase in PDE activity and the calcium dependence of the rate of cGMP turnover (see below); the time scale of adaptation depends largely on the kinetics of this process. The modified cone model below adds a second feedback mechanism (and therefore a second time scale for adaptation), implemented as a calcium-dependent feedback to the cGMP-gated channels; this modification improved fits to long light steps.

In the first step of the model, the stimulus (*Stim*) activates opsin molecules (denoted as *R* for Receptor, *R\** when active), which decay with a rate constant *σ*:

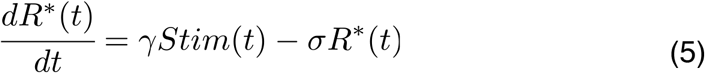

Here, *γ* is a scaling factor (or “opsin gain” factor) that controls the overall sensitivity of the model to light inputs.

Active opsin molecules then activate phosphodiesterase (PDE) molecules through transducin (a delay we assume is negligible) (Pugh & Lamb, 1993), so that the activity of PDE (*P*) follows:

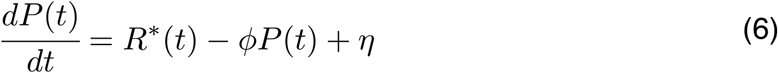

where *ϕ* is the decay rate constant of PDE and *η* is the PDE activity in darkness. The concentration of cGMP in the outer segment (*G*) depends on the PDE-mediated hydrolysis and the rate of synthesis (*S*) by the guanylate cyclase (GC) :

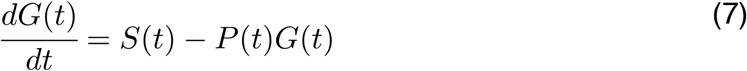

The outer segment current carried by the cGMP-gated channels depends on *G* and has been approximated as:

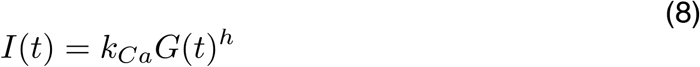

This approximation is valid for current values that are well below the maximal current (Rieke & Baylor, 1996). In this equation *h* denotes the apparent cooperativity and *k*_*Ca*_ is a constant that depends of the maximal current and the affinity of the channel for cGMP. Additionally, we have made this constant calcium-dependent (see below), as a means to introduce feedback to the cGMP-gated channels (Korenbrot, 2012).

A fraction (*q*) of the outer segment current (*I*) is carried by calcium, so that upon exposure to light the calcium concentration (*Ca*) decreases. Calcium extrusion in the outer segment is mainly mediated through the Na^+^/K^+^, Ca^2+^ exchanger. We simplify this process in the model as a single exponential process with time constant *β*:

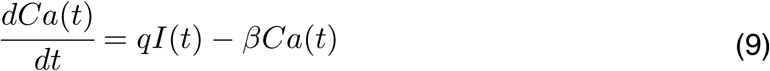

The calcium concentration regulates *S* (the rate of cGMP synthesis) following a Hill curve, such that:

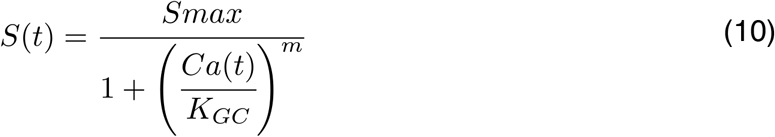

where *S*_*max*_ is the maximum synthesis rate, and *K*_*GC*_ and *m* are the affinity and cooperativity constants respectively.

We also modeled the second feedback as a single-exponential process that is calcium dependent and has a slower decay time constant (*β*_*slow*_):

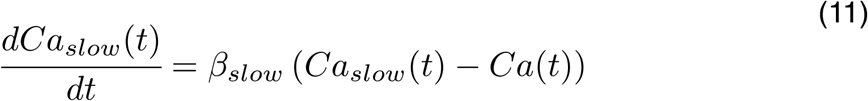

This process determines the value of k_Ca_ (Korenbrot, 2012) such that:

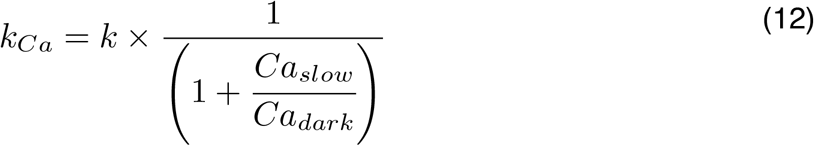

For model fitting, we fixed *k* (0.02 pA.μM^-3^), *h* (3), *m* (4), and *C*_*dark*_ (1 μM) (Robson & Frishman, 1996; Rieke & Baylor, 1996). We then calculated the concentration of cGMP in darkness (*G*_*dark*_) using the measured holding current in darkness (*I*_*dark*_) and equation (8), and the values of *q* and *S*_*max*_ *derived* from steady-state conditions (e.g. in the dark, dCa/dt = 0). We further simplified our model by making *σ* and *ϕ* have equal value, as preliminary fitting showed little advantage to making them differ. We subsequently used a combination of manual trial-and-error and automatic fitting routines in Matlab (*fmincon, lsqcurvefit* and *nlinfit*) to find values for the remaining 6 model parameters (*σ, η, K*_*GC*_, *β, β*_slow_ and *γ*) that would provide a good fit to the response to naturalistic stimuli (Figure 1B) and at the same time accurately predict the single photon-response (Figure 1C). Table 1 shows a summary of model parameters across cones and stimuli.

We evaluated the quality of model fits using the ratio between the sum of absolute errors of the model to the data and the sum of absolute errors of the model to the mean response (i.e. the fraction of variance explained). Model fits were relatively insensitive to modest (∼5%) changes in parameters and were robust to initial conditions for the parameter search. Varying individual parameters by 5% changed the error in the prediction by <10%, with the largest changes associated with manipulations of ϕ or η.

To assess the ability of the model to generalize across cones and stimuli, we allowed *G*_*dark*_ (to match each cell’s *I*_*dark*_) and *γ* (to account for differences in absolute sensitivity between recorded cones) to vary while holding other parameters fixed. Simultaneously fitting responses to a variety of stimuli (naturalistic stimuli (Figure 1B), the single photon response (Figure 1C), steps and flashes (Figure 2B) binary noise and sinusoids (Figure 5C)) or subsets of these stimuli produced model parameters that differed by less than 5% from those fit to the naturalistic stimuli and the single photon response. Including the naturalistic stimuli in the fitting procedure was particularly effective in producing models that generalized to other stimuli. The fraction of variance explained for these different fitting approaches varied minimally (< 5%).

### Alternative models of cone phototransduction

As a first alternative to our phototransduction model, we fit the same dataset to a model that did not include the slower calcium-feedback to the cGMP-gated channel.

This model follows our phototransduction model from equations (5) to (10) and removes the parameter *β*_*slow*._ We followed the same fitting strategy as for the full model. In general, this model behaved well, with similar adaptation values and kinetics, but fits to the naturalistic stimuli or to bright steps suffered because of mismatches in the final currents at the end of fixations (Figure 6 — Supplement 1). As we have shown, we consider this is an important aspect of cone responses, leading us to focus on the biophysical model with two feedback processes.

As a second alternative to our model, we explored an empirical model that is able to capture the responses of turtle cones to a variety of stimuli (Clark et al., 2013). In this model, the light stimulus provides the input to two separate pathways. In the first pathway, the stimulus is directly convolved with a linear filter (*Ky*) before passing through a dynamic low-pass filter that dictates the response of the model. In the second pathway, the stimulus is directly convolved with a slower and delayed linear filter (*Kz*) that dictates the amplitude and time constant of the low-pass filter, providing a way to directly modulate the model’s output (Figure 6 — Supplement 2A). This feedforward implementation of adaptation imparts the model with a mechanism that controls both gain and kinetics in a history-dependent manner. The linear filters are determined by:

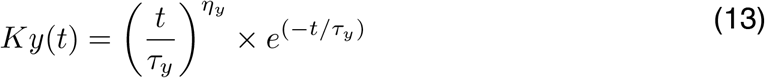

and,

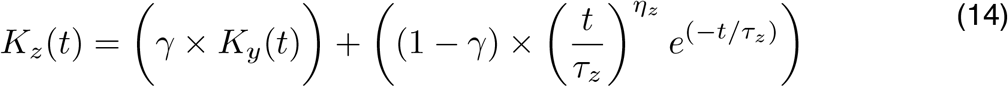

where *τ*_*y*_ and *τ*_*z*_ determine the time-scale of the filters and *η*_*y*_ and *η*_*z*_ determine their rise behavior (Clark et al., 2013). The final output of the model, *r*, is determined by:

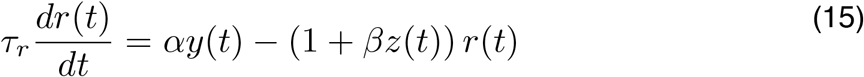

where *α, β*, and *τ*_*r*_ are constants and

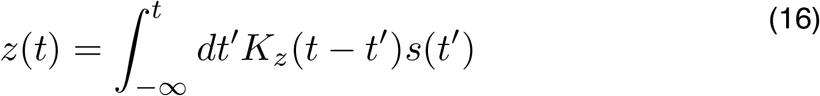

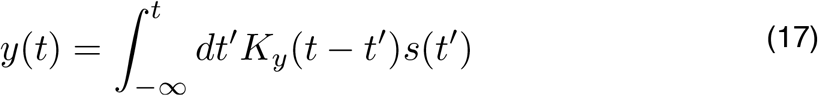

To find fits for this model, we first found a fit to the estimated single-photon response while eliminating adaptation (forcing *β* = 0), allowing us to find values for *τ*_*y*_, *η*_*y*_and *τ*_*r*_ that matched the kinetics of dim-flash responses. After fixing these three values, we fit the response to the naturalistic stimulus to determine values for the other five parameters, namely *τ*_*y*_, *η*_*y*_, *γ, α* and *β* (Figure 6 — Supplement 2B). The fit deviates from the data in two important aspects. First, during transitions from a high to a low light level, the cone response dips below baseline but the model does not. Second, the degree of adaptation required to fit the response to the naturalistic stimulus changes the overall gain of the model such that the model’s single-photon response is ∼10-fold smaller than measured (Figure 6 — Supplement 2C). To test for generalization of this model we fixed the parameters that dictate the kinetics of the model from the fit to the naturalistic stimulus, and let the parameters that dictate gain, sensitivity and strength of feedback (*α, β* and *γ*) to vary while fitting the other data sets. The model is capable of replicating the responses to steps and flashes, with adequate degree and kinetics of gain changes, especially for adaptation onset. These model fits were unable to capture the changes of steady-state current and the dependence of gain on background, requiring ∼300-fold higher light levels for half-adaptation than real cones (Figure 6 — Supplement 2G-J). Model parameters that bridged this discrepancy greatly distorted fits to the other datasets and were not pursued further.

We additionally explored a modification of the empirical model, in which we added a second adaptation mechanism with a longer timescale (Figure 6 - Supplement 3A), such that equation 14 is replaced by:

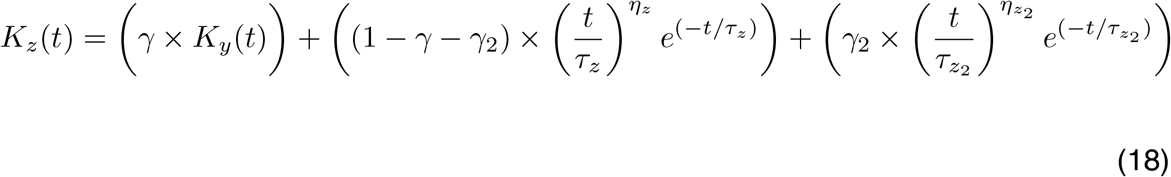

adding three new parameters (*γ*_*z2*_, *τ*_*z2*_and *η*_*z2*_). We followed the same fitting strategy as for the previous empirical model and found that fits improved just slightly and limitations persisted, in particular the inability to capture the dependence of steady state-current and gain on background and the dips of current below baseline after transitions from high to low light levels (Figure 6 -Supplement 3B-J). Preliminary attempts to fix these issues by adding some intrinsic activity to the model (akin to a cone’s dark current) seemed promising but required adding more parameters to the existing eleven, approaching the number of free parameters of our biophysical model.

As a final benchmark, we fit all models to example cone responses to the high-contrast binary noise and sinusoidal stimuli presented in Figure 5. All models were able to adequately capture responses to the both stimuli when directly fit to example traces (Figure 6 - Supplement4A and Supplement5A), and displayed acceptable asymmetric responses when compared to population data (Figure 6 - Supplement4B and Supplement5B). As expected from fits to other datasets, the empirical models recover from adaptation faster than real cones, producing deviations in transitions from high to low light levels, and struggle to capture the responses to sinusoids at the lowest light levels. The phototransduction models fail to capture responses to these stimuli at the highest light levels (near and above 100,000 R*/s), emphasizing that other adaptation mechanisms not included in our model most likely shape responses at high light levels.

### Light-adaptation clamp

Figure 8 uses the biophysical cone model to design stimuli that minimize nonlinearities in the cone responses. We used two models of the cone responses to identify these stimuli: (1) the full biophysical model; (2) a linear model. The linear model was determined by the response of the full model to a brief, low contrast flash (i.e. a flash within the linear range of the full model behavior). The stimulus for the full model was a transformed version of the original stimulus, while the original stimulus (untransformed) provided input to the linear model. We then sought a stimulus transformation that minimized the difference between the outputs of the two models. For sinusoidal stimuli, such as those in Figure 8A, this is particularly simple: the response of the linear model to these stimuli is also sinusoidal, and hence our procedure identifies a stimulus to the full model that creates a sinusoidal output. We refer to this as a “light-adaptation clamp” because the procedure aims to “clamp” cone responses to track a desired response (in this case one that lacks adaptation).

We identified the appropriate stimulus transformation using a gradient-descent approach. We discretized the stimulus into time bins and then perturbed the stimulus at these discrete times. We retained perturbations that decreased the mean-squared difference between the two models’ responses (see Figure 8-Figure Supplement 1 for an example) using Matlab’s fminsearch algorithm. We iterated this process while decreasing the size of the time bins until achieving a stable minimum of the mean-square difference.

---

**Figure 2 - Figure Supplement 1:**
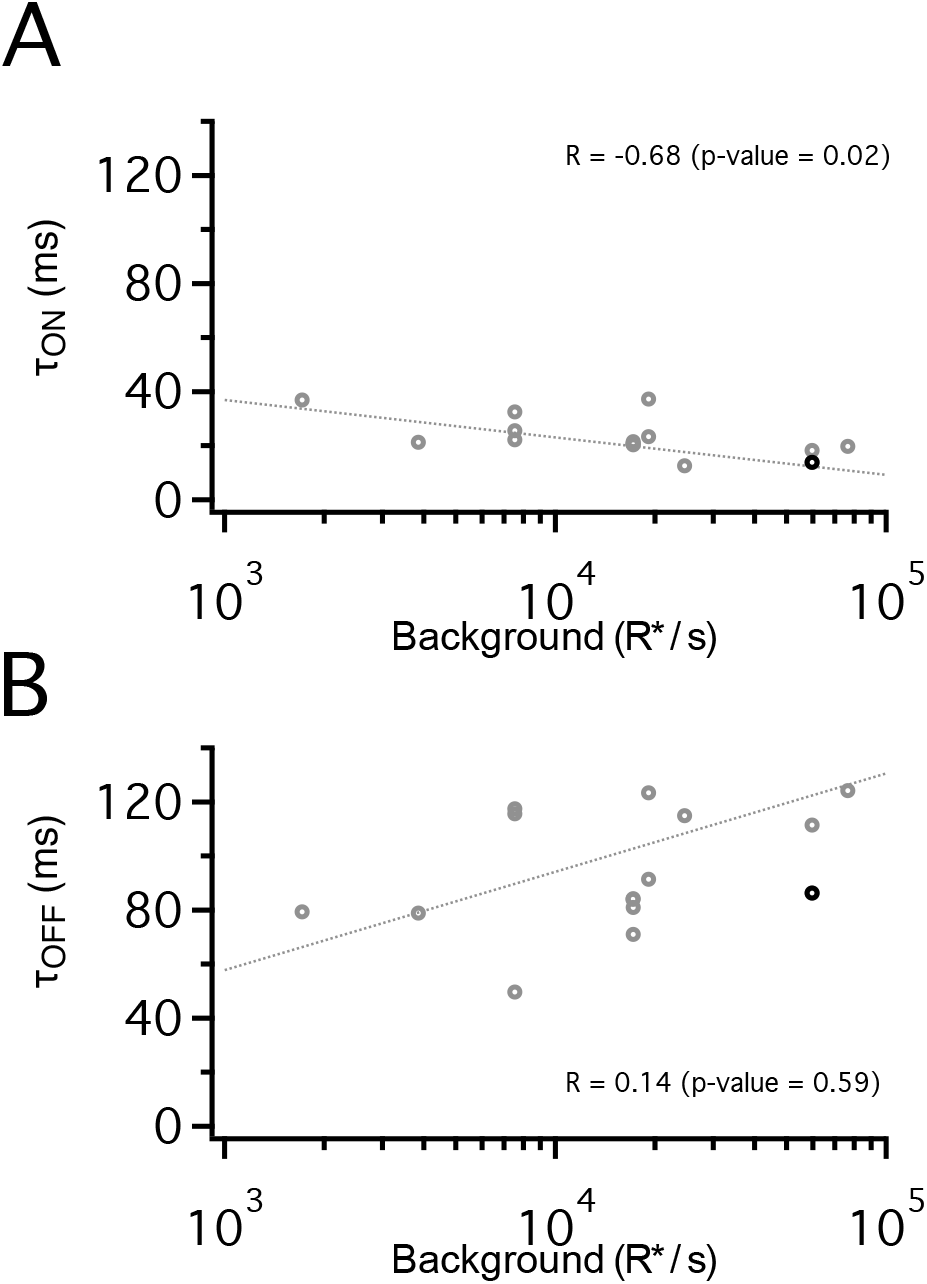
Dependence of adaptation kinetics on light intensity. A. Time constant for onset of adaptation plotted against mean light intensity. B. Time constant of offset of adaptation plotted against mean light intensity. Bold point is example cell from Figure 2.

**Figure 5 - Figure Supplement 1:**
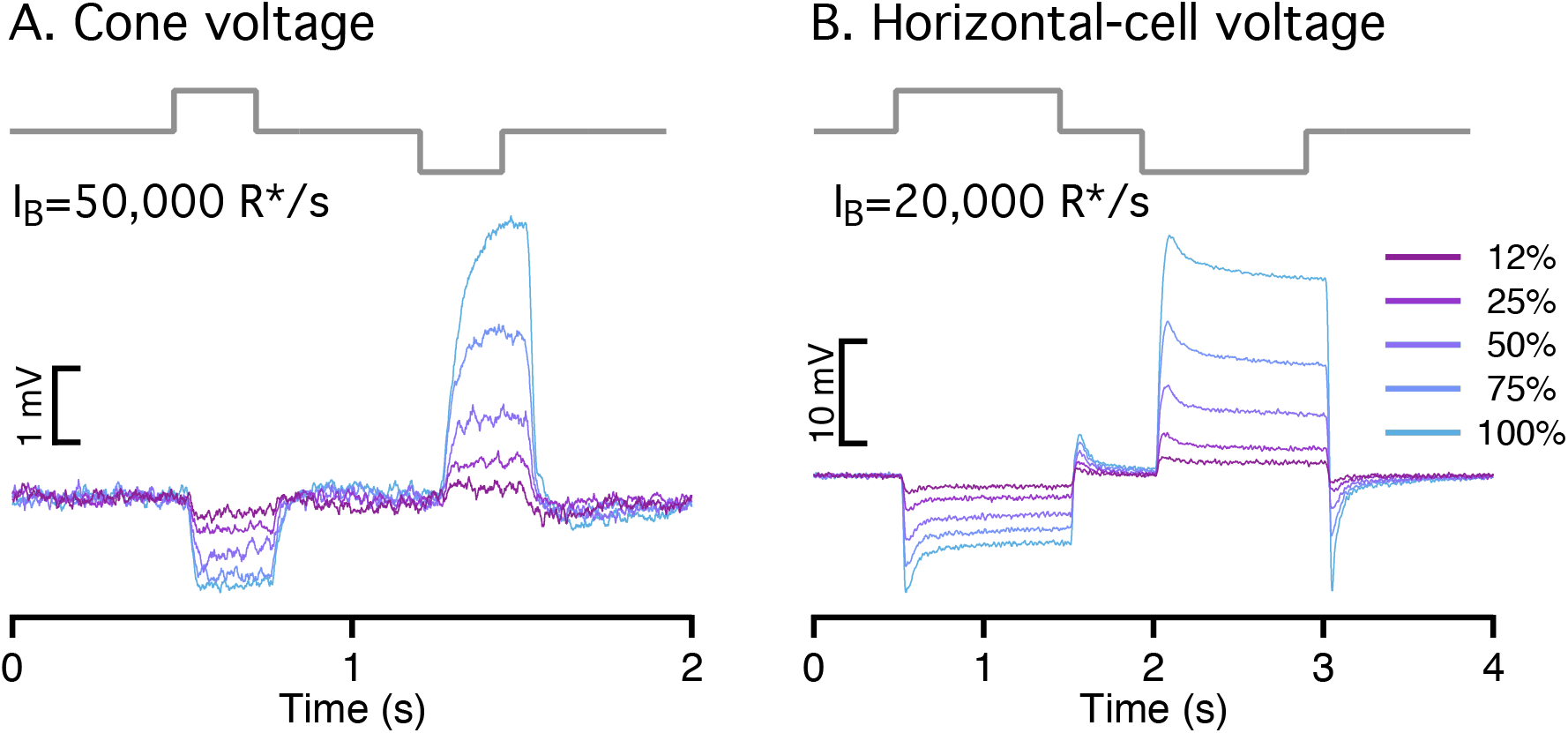
Cone voltages and synaptic output exhibit asymmetric responses to light increments and decrements. A. Cone voltage responses (current-clamp recording) elicited by a family of light increments and decrements. B. Horizontal voltage responses elicited by increments and decrements.

**Figure 6 - Figure Supplement 1:**
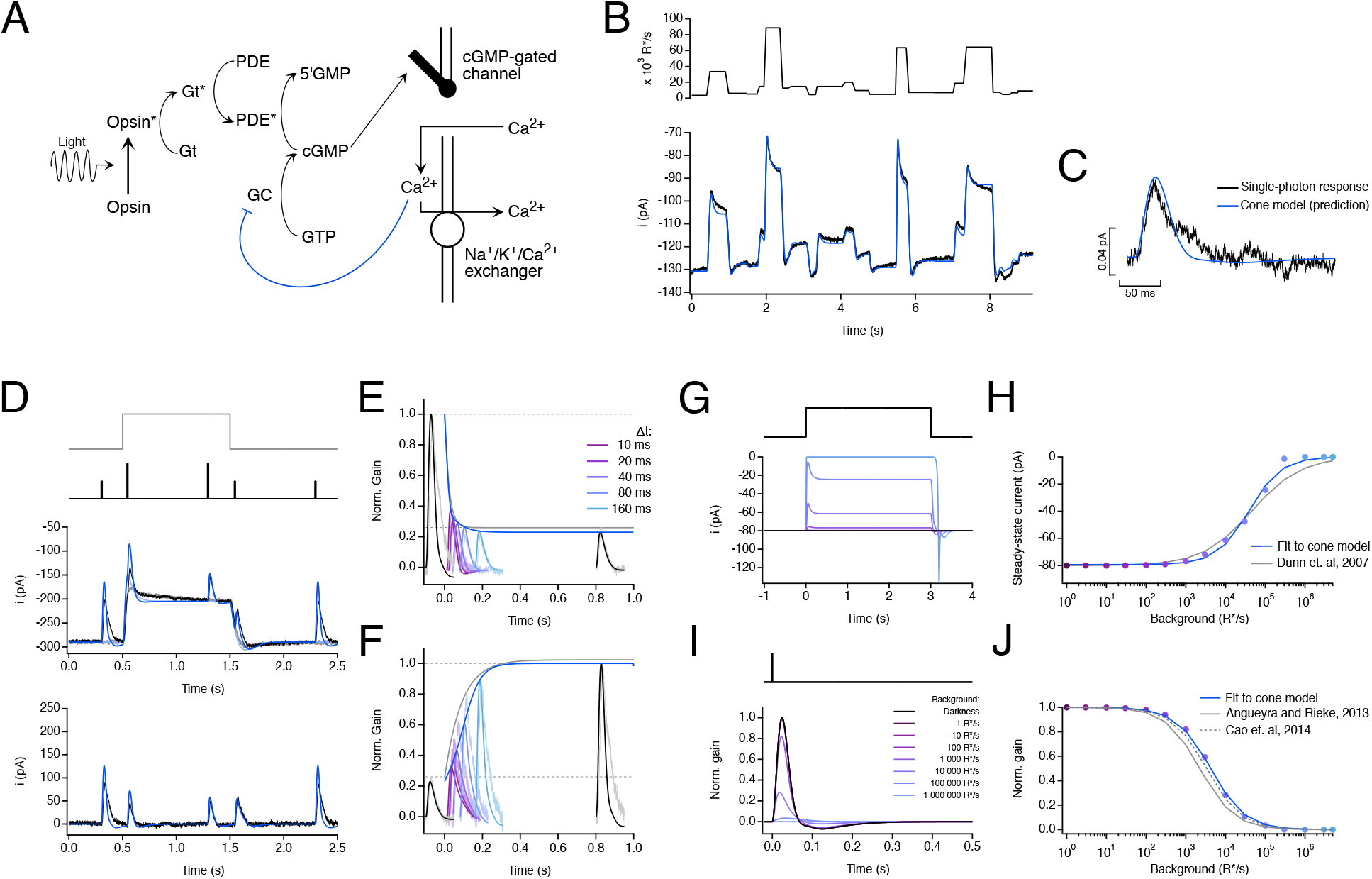
A biophysical model of phototransduction with a single adaptation mechanism performs well but does not capture responses to long steps. A. Schematic of phototransduction cascade and corresponding components of the biophysical model: cyclic-GMP (cGMP) is constantly synthesized by guanylate cyclase (GC), opening cGMP-gated channels in the membrane. Light-activated opsin (Opsin*) leads to channel closure by activating the G-protein transducin (Gt*) which activates phosphodiesterase (PDE*) and decreases the cGMP concentration. Calcium ions (Ca2+) flow into the cone outer segment through the cGMP-gated channels and are extruded through Na+/K+/Ca2+ exchangers in the membrane. Only one feedback mechanism was implemented as a calcium-dependent processes that affect the rate of cGMP-synthesis. B. Fit to cone response to the naturalistic stimulus shown in Figure 1. The model is able to capture the response transients following rapid changes in the stimulus but slightly misses the currents at the end of fixations. See Table 1 - Supplement 1 for fit parameters. C. This model also accurately predicts the amplitude and kinetics of the single-photon response. D-F. Model fit to step and flashes responses from Figure 2. The model exhibits fast changes in gain at step onset (τOn-Model = 22.3 ms) and a slower recovery of gain at step offset (τOff-Model = 122.1 ms). G. Model responses to steps of increasing light intensity. H. Dependence of model’s steady-state current on background light intensity (colored dots). This relation was fit with a Hill equation with a half-maximal background, I_½_ = 38,785 R*/s and a Hill exponent, n = 1.07. I. Estimated single-photon responses of the model, normalized by the response in darkness, at increasing background light intensities. J. Relation of the model’s peak sensitivity, normalized to the peak sensitivity in darkness, across background light-intensity (colored dots). The half-desensitizing background (I_0_) for the model is 4,198 R*/s.

**Figure 6 - Figure Supplement 2:**
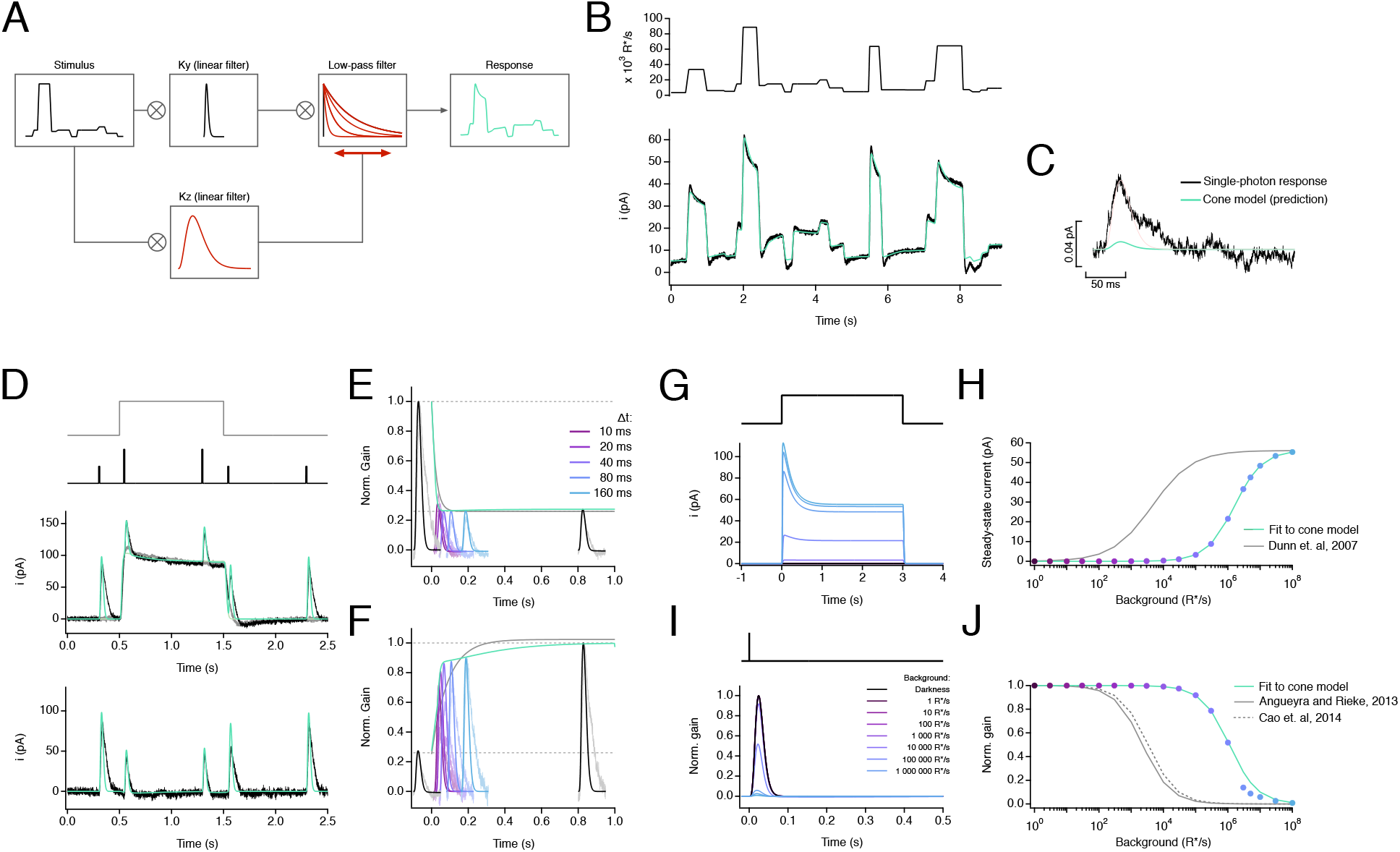
An empirical model of cone responses with a single adaptation mechanism fails to generalize well across cells. A. Schematic of empirical model (Clark et al, 2013) where the stimulus is convolved with a linear filter (Ky) and a dynamic low-pass filter. The time course and amplitude of the low-pass filter are determined by the convolution of the stimulus with a slower linear filter (Kz), which acts as a feed-forward mechanism that dynamically modulates the model’s response. B. Fit to cone response (after baseline subtraction) to the naturalistic stimulus shown in Figure 1. The model is able to capture the response transients following rapid changes in the stimulus but is unable to capture current undershoot in light to dark transitions. See Table 1 - Supplement 2 for fit parameters. C. This model also underestimates the amplitude of the single-photon response by ∼10-fold. D-F. Model fit to step and flashes responses from Figure 2. The model exhibits fast changes in gain both at step onset (τOn-Model = 12.96 ms) and at step offset (τOff-Model = 45.20 ms). G. Model responses to steps of increasing light intensity. H. Dependence of model’s steady-state current on background light intensity (colored dots). This relation was fit with a Hill equation with a half-maximal background, I_½_ = 1,613,990 R*/s and a Hill exponent, n = 1. I. Estimated single-photon responses of the model, normalized by the response in darkness, at increasing background light intensities. J. Relation of the model’s peak sensitivity, normalized to the peak sensitivity in darkness, across background light-intensity (colored dots). The half-desensitizing background (I_0_) for the model is 1,111,790 R*/s.

**Figure 6 - Figure Supplement 3:**
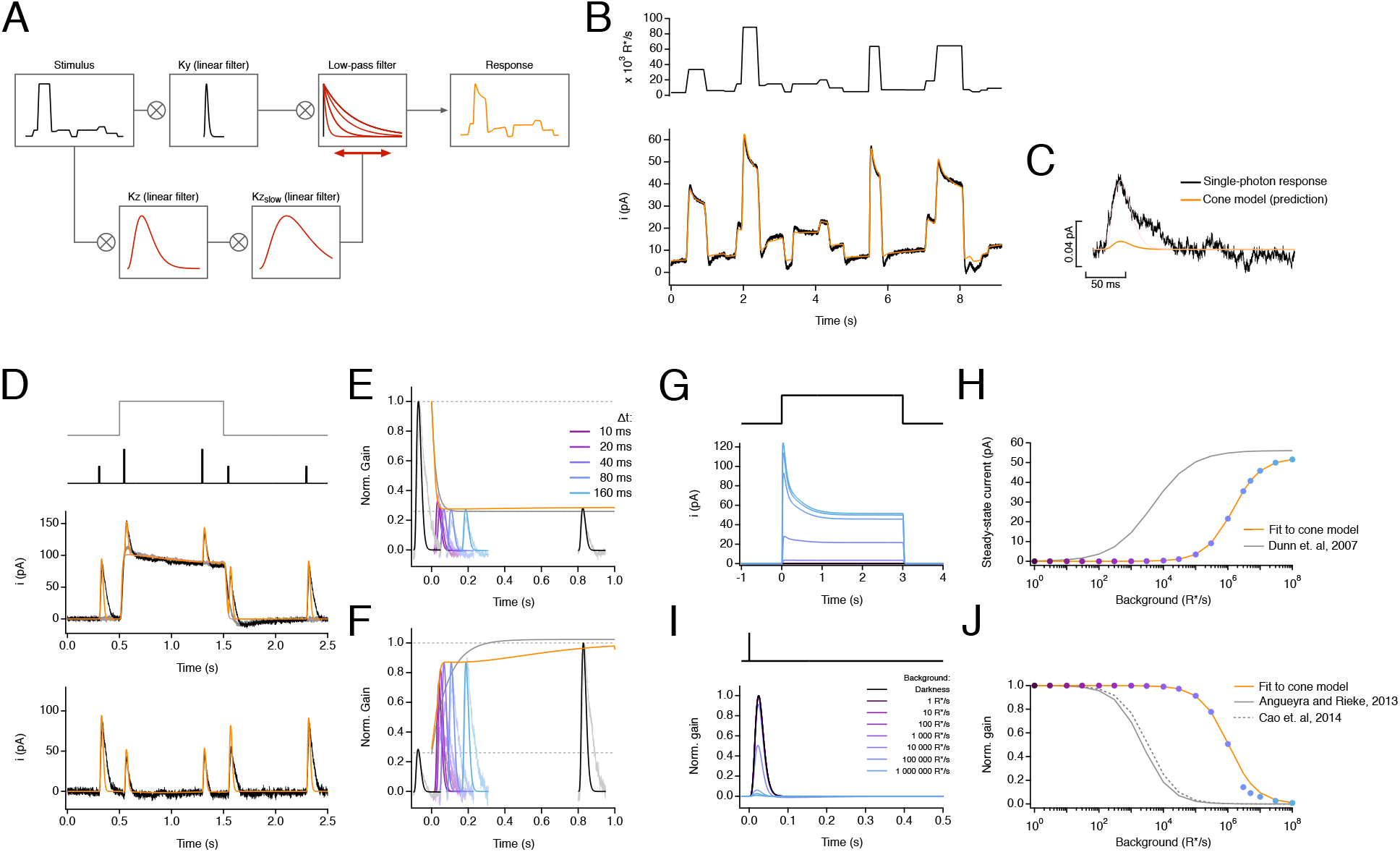
An empirical model of cone responses with a double adaptation mechanism also fails to generalize across cells. A. Schematic of empirical model where the stimulus is convolved with a linear filter (Ky) and a dynamic low-pass filter. The time course and amplitude of the low-pass filter are determined by the successive convolution of the stimulus with two linear filters (Kz and Kz_slow_), providing a feed-forward mechanism that dynamically modulates the model’s response with two different time scales. B. Fit to cone response (after baseline subtraction) to the naturalistic stimulus shown in Figure 1. The model is able to capture the response transients following rapid changes in the stimulus but is still unable to capture current undershoot in light to dark transitions. See Table 1 - Supplement 2 for fit parameters. C. This model also underestimates the amplitude of the single-photon response by ∼10-fold. D-F. Model fit to step and flashes responses from Figure 2. The model exhibits fast changes in gain both at step onset (τOn-Model = 13.38 ms) and at step offset (τOff-Model = 34.75 ms). G. Model responses to steps of increasing light intensity. H. Dependence of model’s steady-state current on background light intensity (colored dots). This relation was fit with a Hill equation with a half-maximal background, I_½_ = 1,428,240 R*/s and a Hill exponent, n = 1. I. Estimated single-photon responses of the model, normalized by the response in darkness, at increasing background light intensities. J. Relation of the model’s peak sensitivity, normalized to the peak sensitivity in darkness, across background light-intensity (colored dots). The half-desensitizing background (I_0_) for the model is 1,052,580 R*/s.

**Figure 6 - Figure Supplement 4:**
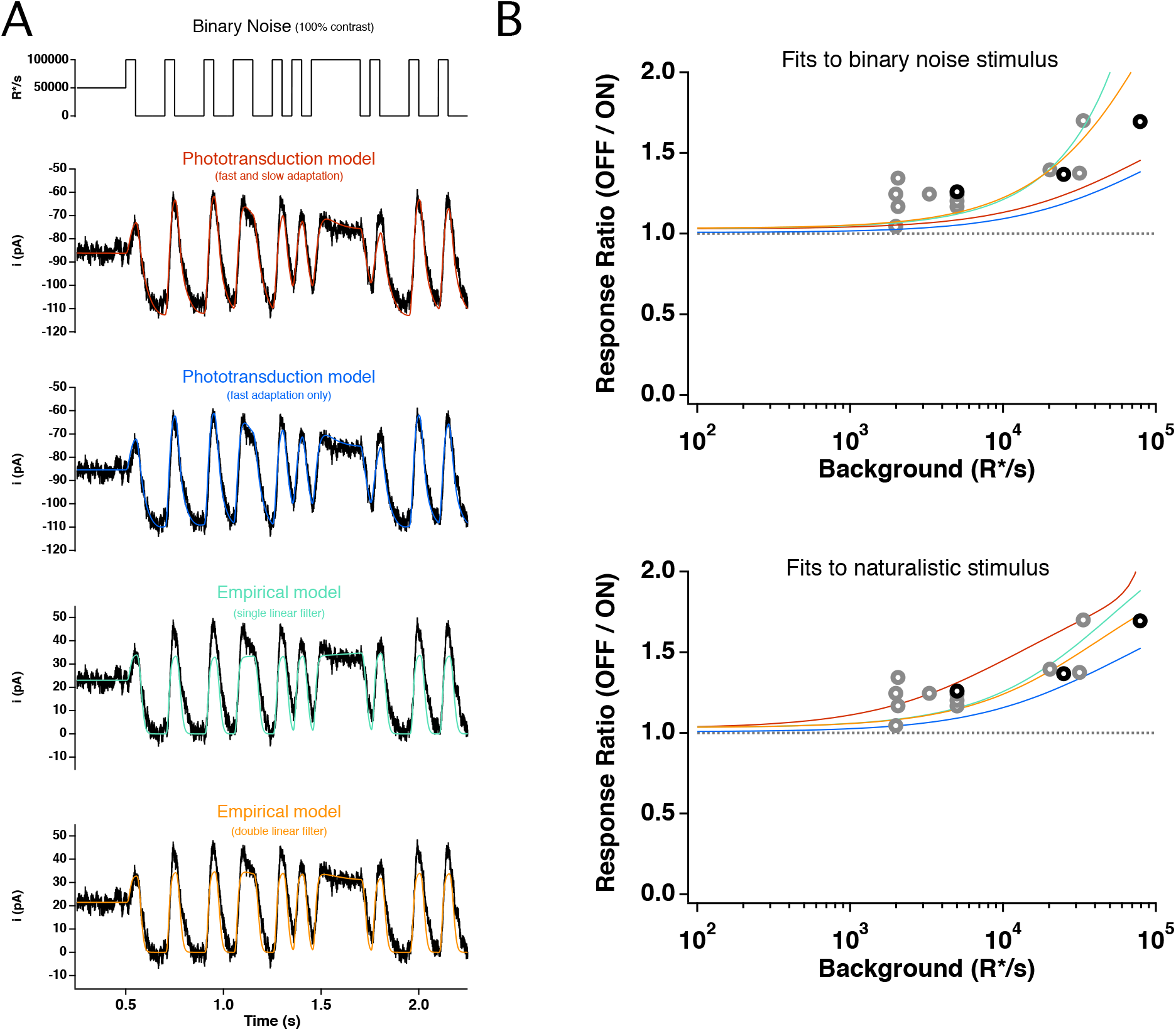
Model responses to binary noise stimuli. A: 100% contrast binary noise stimulus (top trace) and cone photocurrent response (bottom traces), as shown in Figure 5, overlaid with direct fits of each model. B. Ratio of mean negative to mean positive response to binary noise for each model as a function of mean light intensity, derived from fits to example cell in A (top panel) or from fits to naturalistic stimulus, as shown in Figure 1. All models are able to adequately capture the asymmetric responses to this stimulus.

**Figure 6 - Figure Supplement 5:**
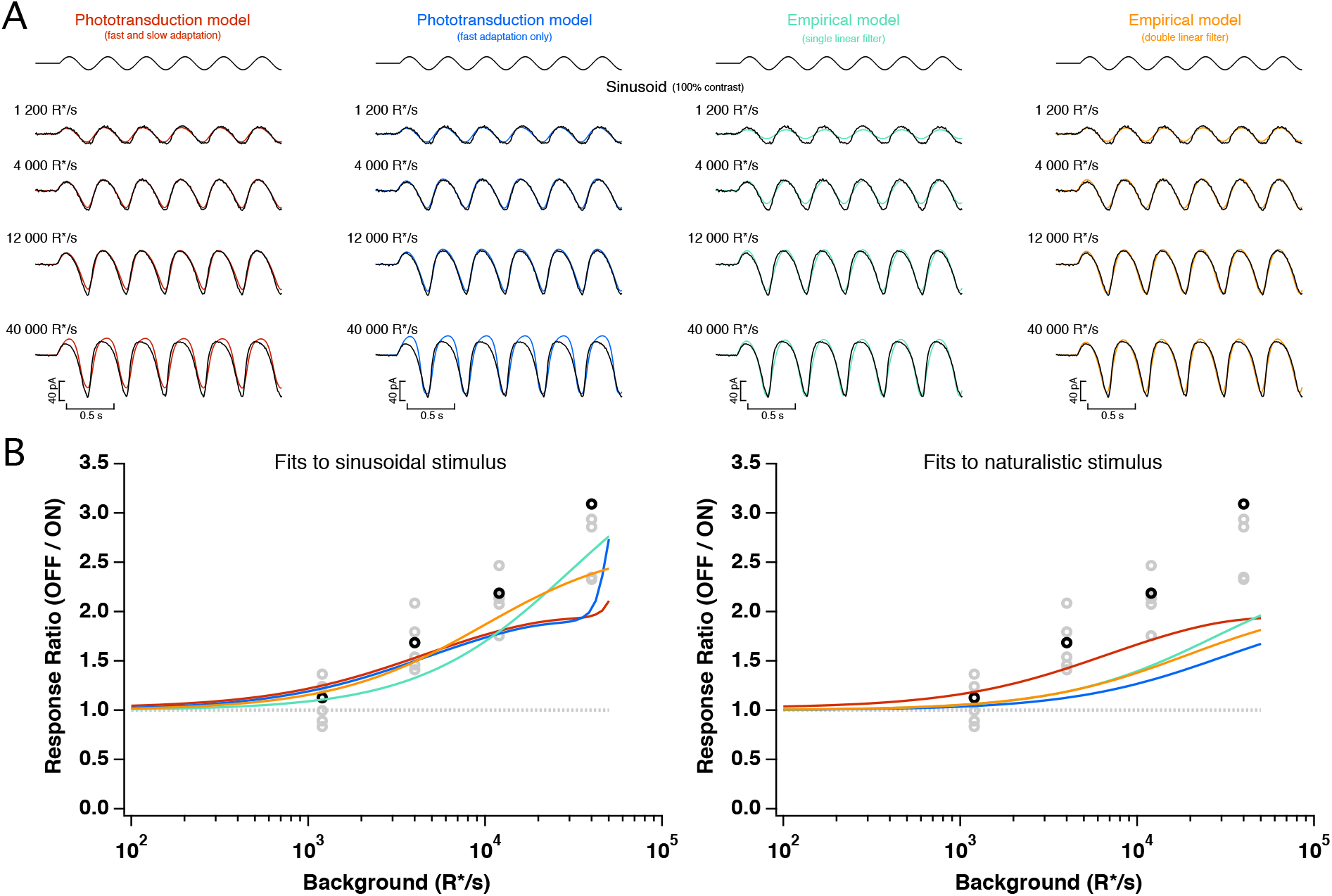
Model responses to sinusoidal stimuli. A: 100% contrast sinusoidal stimulus (top traces) and cone photocurrent response (bottom traces), as shown in Figure 5, overlaid with direct fits of each model. B. Ratio of peak negative to peak positive response to the sinusoidal stimulus for each model as a function of mean light intensity, derived from fits to the example cell in A (top panel) or from fits to naturalistic stimulus, as shown in Figure 1. All models are able to capture the asymmetric responses to this stimulus.

**Figure 7 - Figure Supplement 1:**
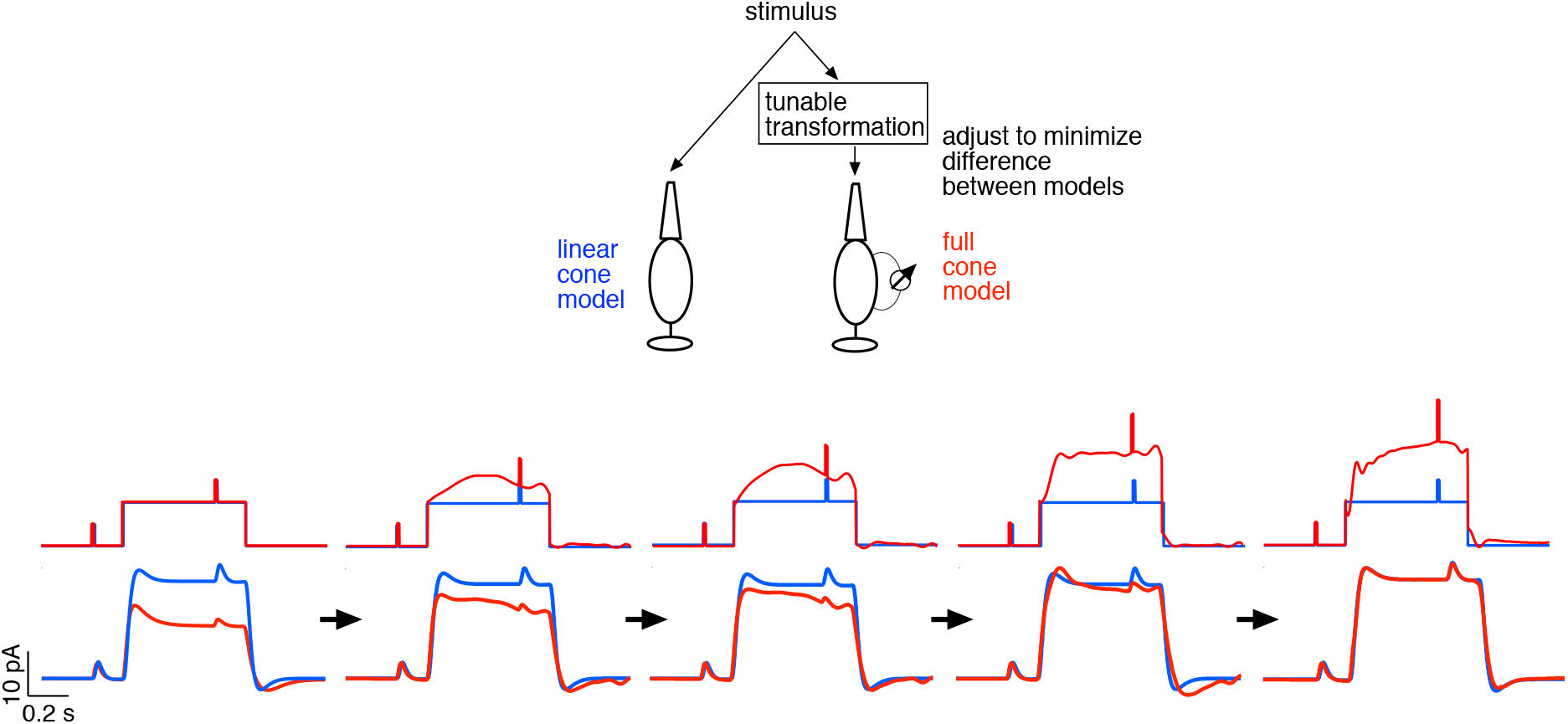
Example of cone light-adaptation clamp procedure. The top panel illustrates the approach. The stimulus to the linear cone model is held fixed, while the stimulus to the full cone model is adjusted until the two models produce similar outputs. The bottom panels show this process for a step and flashes stimulus. Initially (far left) the two stimuli are identical and the two models produce very different outputs, due to adaptation in the full model. Moving rightwards, each panel shows a step in the transformation process, with the final result in the far right.

**Supplement Table 1.**
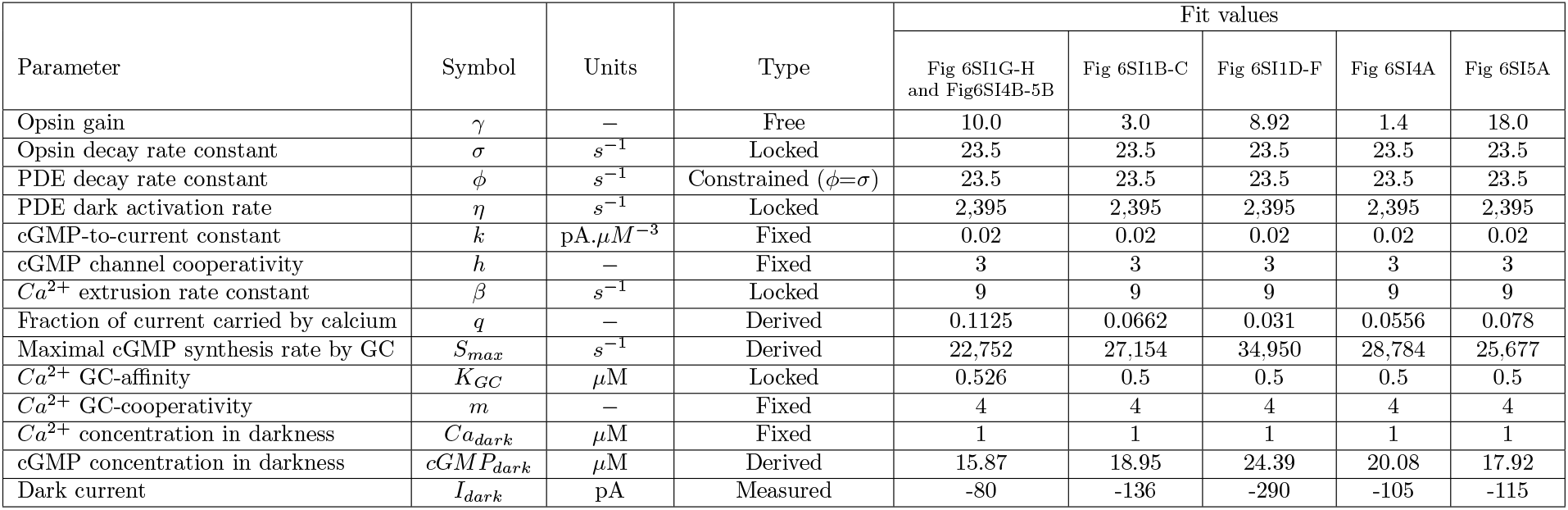
Parameters and best fit values for cone-phototransduction biophysical model with fast adaptation only

| Parameter | Symbol | Units | Type | Fit values |  |  |  |  |
| --- | --- | --- | --- | --- | --- | --- | --- | --- |
|  |  |  |  | Fig 6SI1G-H<br>and Fig6SI4B-5B | Fig 6SI1B-C | Fig 6SI1D-F | Fig 6SI4A | Fig 6SI5A |
| Opsin gain | $\gamma$ | — | Free | 10.0 | 3.0 | 8.92 | 1.4 | 18.0 |
| Opsin decay rate constant | $\sigma$ | $s^{-1}$ | Locked | 23.5 | 23.5 | 23.5 | 23.5 | 23.5 |
| PDE decay rate constant | $\phi$ | $s^{-1}$ | Constrained ($\phi=\sigma$) | 23.5 | 23.5 | 23.5 | 23.5 | 23.5 |
| PDE dark activation rate | $\eta$ | $s^{-1}$ | Locked | 2,395 | 2,395 | 2,395 | 2,395 | 2,395 |
| cGMP-to-current constant | $k$ | $pA \cdot \mu M^{-3}$ | Fixed | 0.02 | 0.02 | 0.02 | 0.02 | 0.02 |
| cGMP channel cooperativity | $h$ | — | Fixed | 3 | 3 | 3 | 3 | 3 |
| $Ca^{2+}$ extrusion rate constant | $\beta$ | $s^{-1}$ | Locked | 9 | 9 | 9 | 9 | 9 |
| Fraction of current carried by calcium | $q$ | — | Derived | 0.1125 | 0.0662 | 0.031 | 0.0556 | 0.078 |
| Maximal cGMP synthesis rate by GC | $S_{max}$ | $s^{-1}$ | Derived | 22,752 | 27,154 | 34,950 | 28,784 | 25,677 |
| $Ca^{2+}$ GC-affinity | $K_{GC}$ | $\mu M$ | Locked | 0.526 | 0.5 | 0.5 | 0.5 | 0.5 |
| $Ca^{2+}$ GC-cooperativity | $m$ | — | Fixed | 4 | 4 | 4 | 4 | 4 |
| $Ca^{2+}$ concentration in darkness | $Ca_{dark}$ | $\mu M$ | Fixed | 1 | 1 | 1 | 1 | 1 |
| cGMP concentration in darkness | $cGMP_{dark}$ | $\mu M$ | Derived | 15.87 | 18.95 | 24.39 | 20.08 | 17.92 |
| Dark current | $I_{dark}$ | pA | Measured | -80 | -136 | -290 | -105 | -115 |

**Supplement Table 2.** Parameters and best fit values for empirical model with a single linear filter

| Parameter | Symbol | Units | Type | Fit values |  |  |  |
| --- | --- | --- | --- | --- | --- | --- | --- |
|  |  |  |  | Fig 6SI2B-C,G-H<br>and Fig6SI4B-5B | Fig 6SI2D-F | Fig 6SI4A | Fig 6SI5A |
| alpha | $\alpha$ | — | Free | 19.40 | 1320 | 70 | 350 |
| beta | $\beta$ | — | Free | 0.36 | 8.29 | 0.79 | 0.45 |
| gamma | $\gamma$ | — | Free | 0.448 | 0.745 | 1.370 | 0.600 |
| Ky filter time constant | $\tau_y$ | ms | Locked | 4.48 | 4.48 | 4.48 | 4.48 |
| Ky filter rise constant | $\eta_y$ | — | Locked | 4.33 | 4.33 | 4.33 | 4.33 |
| Kz filter time constant | $\tau_z$ | ms | Locked | 166 | 166 | 166 | 166 |
| Kz filter rise constant | $\eta_z$ | — | Locked | 1.00 | 1.00 | 1.00 | 1.00 |
| Response time constant | $\tau_r$ | ms | Locked | 4.78 | 4.78 | 4.78 | 4.78 |

**Supplement Table 3.** Parameters and best fit values for empirical model with a double linear filter

| Parameter | Symbol | Units | Type | Fit values |  |  |  |
| --- | --- | --- | --- | --- | --- | --- | --- |
|  |  |  |  | Fig 6SI3B-C,G-H<br>and Fig6SI4B-5B | Fig 6SI3D-F | Fig 6SI4A | Fig 6SI5A |
| alpha | $\alpha$ | — | Free | 20.50 | 1248 | 880 | 600 |
| beta | $\beta$ | — | Free | 0.312 | 6.51 | 17.0 | 8.00 |
| gamma | $\gamma$ | — | Free | 0.5 | 0.896 | 0.721 | 0.591 |
| Ky filter time constant | $\tau_y$ | ms | Locked | 4.48 | 4.48 | 4.48 | 4.48 |
| Ky filter rise constant | $\eta_y$ | — | Locked | 4.33 | 4.33 | 4.33 | 4.33 |
| Kz filter time constant | $\tau_z$ | ms | Locked | 35 | 35 | 35 | 35 |
| Kz filter rise constant | $\eta_z$ | — | Locked | 2.84 | 2.84 | 2.84 | 2.84 |
| gamma<sub>2</sub> | $\gamma_2$ | — | Free | 0.146 | 0.114 | 0.06 | 0.01 |
| Kz<sub>2</sub> filter time constant | $\tau_{z2}$ | ms | Locked | 184.0 | 184.0 | 184.0 | 184.0 |
| Kz<sub>2</sub> filter rise constant | $\eta_{z2}$ | — | Locked | 2.32 | 2.32 | 2.32 | 2.32 |
| Response time constant | $\tau_r$ | ms | Locked | 4.78 | 4.78 | 4.78 | 4.78 |

